# Transcriptome changes in chlorsulfuron-treated plants are caused by acetolactate synthase inhibition and not induction of a herbicide detoxification system in *Marchantia polymorpha*

**DOI:** 10.1101/2022.08.31.505973

**Authors:** Alexandra Casey, Thomas Köcher, Samuel Caygill, Clément Champion, Clémence Bonnot, Liam Dolan

## Abstract

A sensing mechanism in mammals perceives xenobiotics and induces the transcription of genes encoding proteins that detoxify these molecules. However, it is unclear if plants sense xenobiotics, and activate an analogous signalling system leading to their detoxification. Using the liverwort *Marchantia polymorpha*, we tested the hypothesis that there is a sensing system in plants that perceives herbicides resulting in the increased transcription of genes encoding proteins that detoxify these herbicides. Consistent with the hypothesis, we show that chlorsulfuron-treatment induces changes in the *M. polymorpha* transcriptome. However, these transcriptome changes do not occur in chlorsulfuron (CS)-treated target site resistant mutants, where the gene encoding the target carries a mutation that confers resistance to chlorsulfuron. Instead, we show that inactivation of the chlorsulfuron target, acetolactate synthase (ALS) (also known as acetohydroxyacid synthase (AHAS)), is required for the transcriptome response. These data are consistent with the changes in the transcriptome of chlorsulfuron-treated plants being caused by disrupted amino acid synthesis and metabolism resulting from acetolactate synthase inhibition. These conclusions suggest that chlorsulfuron is not sensed in *M. polymorpha* leading to induce a detoxification system.

**Author Summary:** Herbicide use is increasing throughout the world, however we know little about how plants respond to herbicide treatment and regulate their metabolism. Some plants have evolved resistance to herbicides such as chlorsulfuron by increasing the detoxification of the herbicide compared to sensitive plants. It has been suggested that plants can directly sense the herbicide chemical which activates a detoxification response, in a similar way to the detoxification of foreign chemicals in mammalian cells. The liverwort *Marchantia polymorpha* is an excellent system to study plant herbicide responses due to its short generation time, ease of propagation and low genetic redundancy. We show that chlorsulfuron treatment alters the expression of many genes in *M. polymorpha*, however plants with a resistance-conferring mutation in the molecular target of chlorsulfuron do not show any changes in gene expression in response to chlorsulfuron treatment. This result indicates that transcriptome changes caused by chlorsulfuron depend on the inhibition of the target by chlorsulfuron. This suggests that plants do not sense chlorsulfuron and activate a detoxification system. This finding has implications for herbicide use and discovery.

## Introduction

In mammals, sensing mechanisms detect the presence of xenobiotics which in turn activate a signalling system that induces the expression of genes that encode proteins that function in their metabolism (1,2). Genes induced by exposure to xenobiotics include those encoding cytochrome P450 monooxygenases that oxidize substrates, making them more polar, which in the case of xenobiotics can make them available for metabolism and inactivation (3). Genes encoding cytochrome P450 monooxygenases are also transcriptionally induced – along with many other genes – in plants treated with herbicides (4–19). Furthermore, transcriptional changes of genes encoding cytochrome P450 monooxygenases have also been shown to confer resistance to herbicides in herbicide-resistant weeds (14,20–25). Consequently, it has been hypothesized that a sensing mechanism exists in plants that induces the transcription of genes encoding enzymes that detoxify herbicides (26,27). An alternative hypothesis is that the detoxification of herbicides in plants is not activated by a herbicide sensing mechanism.

Chlorsulfuron is a member of the hydroxyurea family of herbicides that are active against a wide range of weeds including members of the *Poaceae* (grasses) and eudicots (broad leaf plants). Its target is the enzyme acetolactate synthase (ALS) which functions in the first committed step in the branch chain amino acid pathway (28,29). ALS catalyses the reaction between acetate and pyruvate to produce 2-acetolactate a precursor of valine and leucine. It also catalyses the reaction between acetate and 2-hydroxybutyrate to form 2-aceto-2-hydroxybutyrate, a precursor of leucine. Treatment of plants with chlorsulfuron and other ALS inhibitors results in a reduction in levels of branched chain amino acids in a variety of plants (30–36). However, it is unclear what causes plant death following treatment with ALS inhibitors. It is likely that reduced levels of branched chain amino acids contribute, but it is possible that secondary catastrophic effects that result from ALS inhibition are responsible for death.

Resistance to chlorsulfuron has evolved in the field in response to the intense selection pressure imposed by herbicide treatment (37–39). Two types of resistance have evolved to chlorsulfuron; target site resistance (TSR) and non-target site resistance (NTSR). Target site resistance results from mutations that cause amino acid substitutions in the channel of the ALS protein leading to the active site (40). It has been shown by solving the crystal structure of the *A. thaliana* ALS enzyme in complex with ALS-inhibiting herbicides that these mutations change the shape of the channel and block access of the herbicide to the active site (40). Codons of several amino acids in the channel region mutate to cause TSR (37). For example, a resistance-conferring mutation that is frequently found in resistant weeds is at the codon encoding a conserved proline (P197) (41–43). Non-target site resistance has evolved in weeds in the field and has been shown to result from higher levels of expression of genes encoding enzymes such as cytochrome P450s (14,44). It is hypothesized that the high levels of expression of these enzymes can chemically modify the herbicide making it inactive and or more susceptible to degradation than in wild-type.

To determine if plants sense herbicides we characterized the response of the model plant *M. polymorpha* to chlorsulfuron treatment. Our data are consistent with the hypothesis that chlorsulfuron acts by inhibiting acetolactate synthase. We show chlorsulfuron-treatment changes the transcriptome. However, we demonstrate that the transcriptome changes caused by chlorsulfuron-treatment are not induced in plants harbouring a mutant form of ALS that cannot bind to chlorsulfuron. Therefore, the presence of chlorsulfuron alone is not sufficient to induce the transcriptome change. We predict transcriptome changes induced by chlorsulfuron are not involved in detoxification but result from the physiological changes to the plant caused by the inhibition of the chlorsulfuron target. We test this hypothesis and demonstrate that ectopic overexpression of three cytochrome P450 monooxygenase and glutathione transferase encoding genes do not increase chlorsulfuron resistance. These data suggest that a xenobiotic-sensing mechanism – analogous to the sensing system in mammals – that senses chlorsulfuron, does not exist in *M. polymorpha*.

## Results

### Chlorsulfuron represses the growth of *Marchantia polymorpha*

Since some herbicides do not effectively control *M. polymorpha*, we first tested if chlorsulfuron was potent against this liverwort (45,46). To determine if chlorsulfuron (CS) represses the growth of *M. polymorpha*, Tak-2 (wild-type female) gemmae were plated on solid nutrient media for 7 days and transferred to media containing different concentrations of chlorsulfuron and grown in light for a further 7 days. To quantify gemmaling growth, thallus area was calculated when imaged from above. The data indicate that chlorsulfuron represses the growth of *M. polymorpha* in a dose dependent manner. The concentration at which plant growth is inhibited by 50% (GR_50_) was 3.3 nM (SD = 0.5) and the lethal dose was estimated to be 300 nM (Fig 1A). These data demonstrate that chlorsulfuron has herbicidal activity on *M. polymorpha* and represses growth in the nM concentration range.

**Fig 1.**
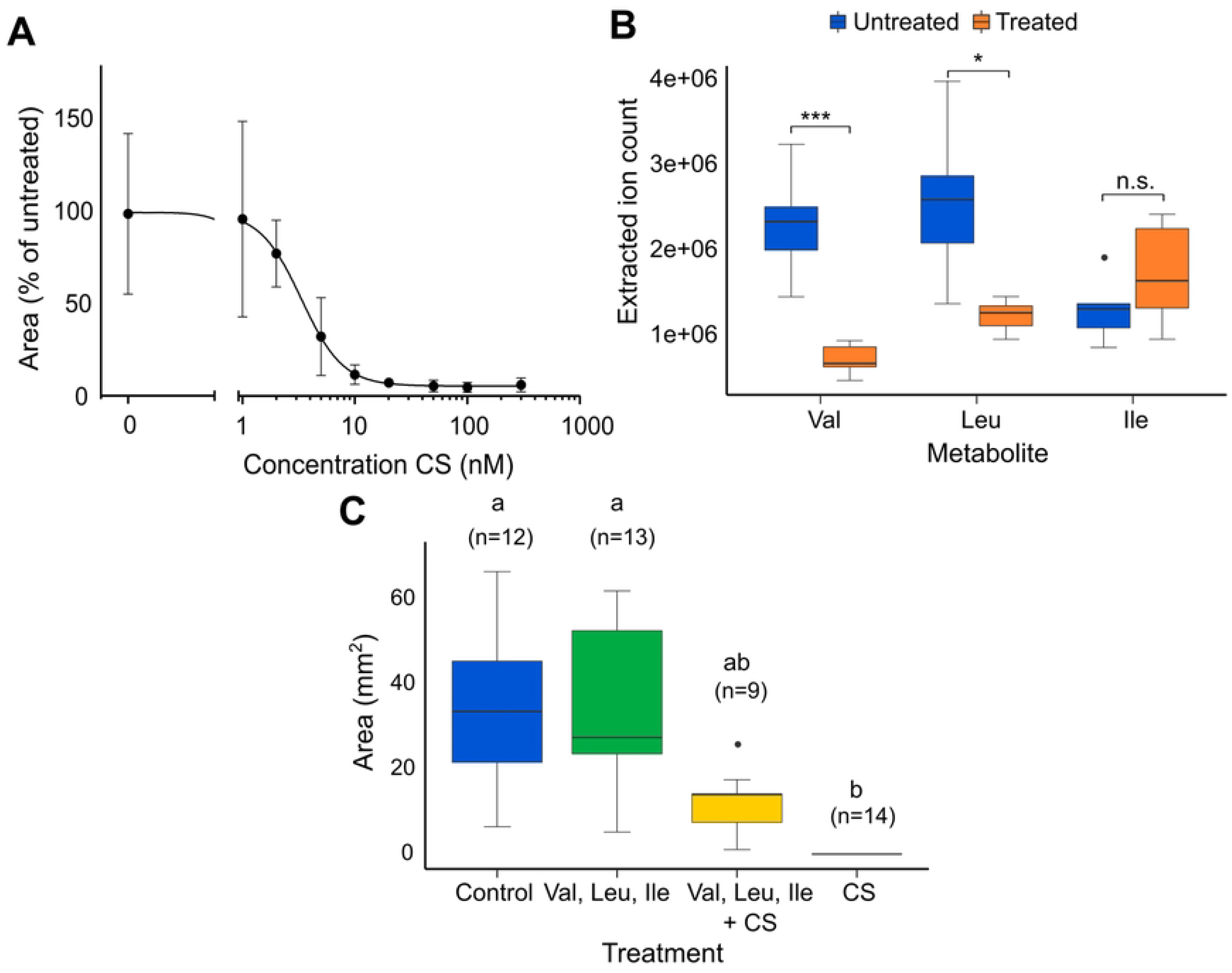
Chlorsulfuron represses the growth of *Marchantia polymorpha*. **(A)** Dose response curve of the growth of 14-day old Tak-2 gemmae after seven days on CS-treated medium. Data plotted with a non-linear fit, sigmoidal, four-parameter regression curve. N=5-10. Error bars are ± SD. **(B)** Targeted metabolic analysis showing the accumulation of valine, leucine and isoleucine, tryptophan, and isopropylmalate in untreated Tak-2 plants and in plants treated with 100 nM CS. Plants were grown for 11 days, transferred to treatment plates for 24 h, then extracted. Extracts were analysed using a targeted approach by LC-MS/MS. Statistical significance based on Student’s t-test between treatment groups for each metabolite. *=p < 0.05, ** = p < 0.01, *** = p < 0.001, n.s. = not significant. N=6 except in isoleucine untreated group n=5 (one outlier was removed). Box and whisker plot where whiskers are first and fourth quartiles, boxes represent second and third quartiles, and the horizontal line is the median. **(C)** 14-day old Tak-2 gemmae grown from day 0 on medium supplemented with 800 µM branched chain amino acids and/or 100 nM CS then imaged using NightOwl imaging system which measures chlorophyll autofluorescence. Plant area (mm^2^) was measured using indiGo software. Statistical significance based on Kruskal-Wallis test, with groups determined by Nemenyi post-hoc test. Box and whisker plot where whiskers are first and fourth quartiles, boxes represent second and third quartiles, and the horizontal line is the mean.

Inhibition of ALS by chlorsulfuron has been shown to block the biosynthesis of branched chain amino acids in several organisms including bacteria and flowering plants (30–36). To test the hypothesis that chlorsulfuron has the same mode of action in *M. polymorpha* as in other organisms, we determined the impact of chlorsulfuron-treatment on the levels of branched chain amino acids. Plants were grown from gemmae for 11 days in constant light and then transferred to solid media containing 100 nM chlorsulfuron or untreated media for 24 hours. Plants were harvested and small molecules were extracted and separated by liquid chromatography before detection with mass spectroscopy. The levels of the branched chain amino acids valine and leucine were significantly lower in treated plants than in untreated controls. Valine accumulated to approximately 725,000 ions (SD = 180,000) in chlorsulfuron-treated plants compared to untreated control where the amino acid accumulated to 2,310,000 ions (SD = 600,000) (Fig 1B). Leucine accumulated to 1,240,000 ions (SD = 190,000) in chlorsulfuron-treated plants compared to untreated control where the amino acid accumulated to 2,580,000 ions (SD = 890,000). There was no significant difference in the amount of isoleucine between the chlorsulfuron-treated and untreated plants. These data are consistent with the hypothesis that chlorsulfuron blocks the activity of the ALS enzyme, the first committed step in branched chain amino acid synthesis, leading to a reduction of branched chain amino acids in the plant.

To independently test if chlorsulfuron acts by repressing branched chain amino acid synthesis, we tested if the lethality caused by chlorsulfuron-treatment could be suppressed by growing plants in media supplemented with branched chain amino acids. Growing plants in 100 nM chlorsulfuron entirely inhibited growth while untreated controls grew to a mean area of 33.5 (SD = 18.4) mm^2^ (Fig 1C). Growing plants in the presence of 100 nM chlorsulfuron and supplemented with 800 µM valine, leucine and isoleucine restored growth to 12.6 (SD = 7.2) mm^2^. The suppression of the chlorsulfuron-inhibition of growth by branched chain amino acid supplementation is consistent with the hypothesis that chlorsulfuron-treatment blocks the synthesis of branched chain amino acids by inhibiting ALS activity. These data suggest that that the inhibition of growth by chlorsulfuron is, at least in part, the result of a depletion in the pool of branched chain amino acids in chlorsulfuron-treated plants.

Together these data are consistent with the hypothesis that chlorsulfuron inhibits *M. polymorpha* growth by inhibiting branched chain amino acid biosynthesis as it does in other organisms.

### The *ALS* P197 mutation confers resistance to chlorsulfuron

If ALS is the target of chlorsulfuron in *M. polymorpha*, we hypothesized that mutations in codons that encode amino acids important to chlorsulfuron binding could confer resistance to the herbicide. To identify mutations that confer resistance to chlorsulfuron we mutated 10^6^ spores with UV radiation, grew the spores on chlorsulfuron, and selected resistant mutants after 14 days of growth in light. We selected two independent mutants that were morphologically similar to wild-type and verified their resistance by growing gemmae from each on 140 nM chlorsulfuron; both mutants grew while wild-type plants died.

Once chlorsulfuron-resistance was verified, we isolated DNA from each of the two lines and sequenced the Mp*ALS* gene using Sanger sequencing. There was a mutation at the codon for a proline at amino acid position 197 of *A. thaliana* ALS (position 207 of *M. polymorpha* ALS protein sequence) in each mutant that resulted in a P197L change in each (Fig 2A, B). P197 was unchanged in wild-type parental lines. The presence of the P197L mutations in Mp*ALS* in both chlorsulfuron-resistant mutants is consistent with the hypothesis that chlorsulfuron acts by binding to Mp*ALS*. Mutants were designated Mp*acetolactate synthase*^*chlorsulfuronresistantP197L*^ (Mp*als*^*csP197L_7*^) and Mp*als*^*csP197L_10*^ respectively.

**Fig 2.**
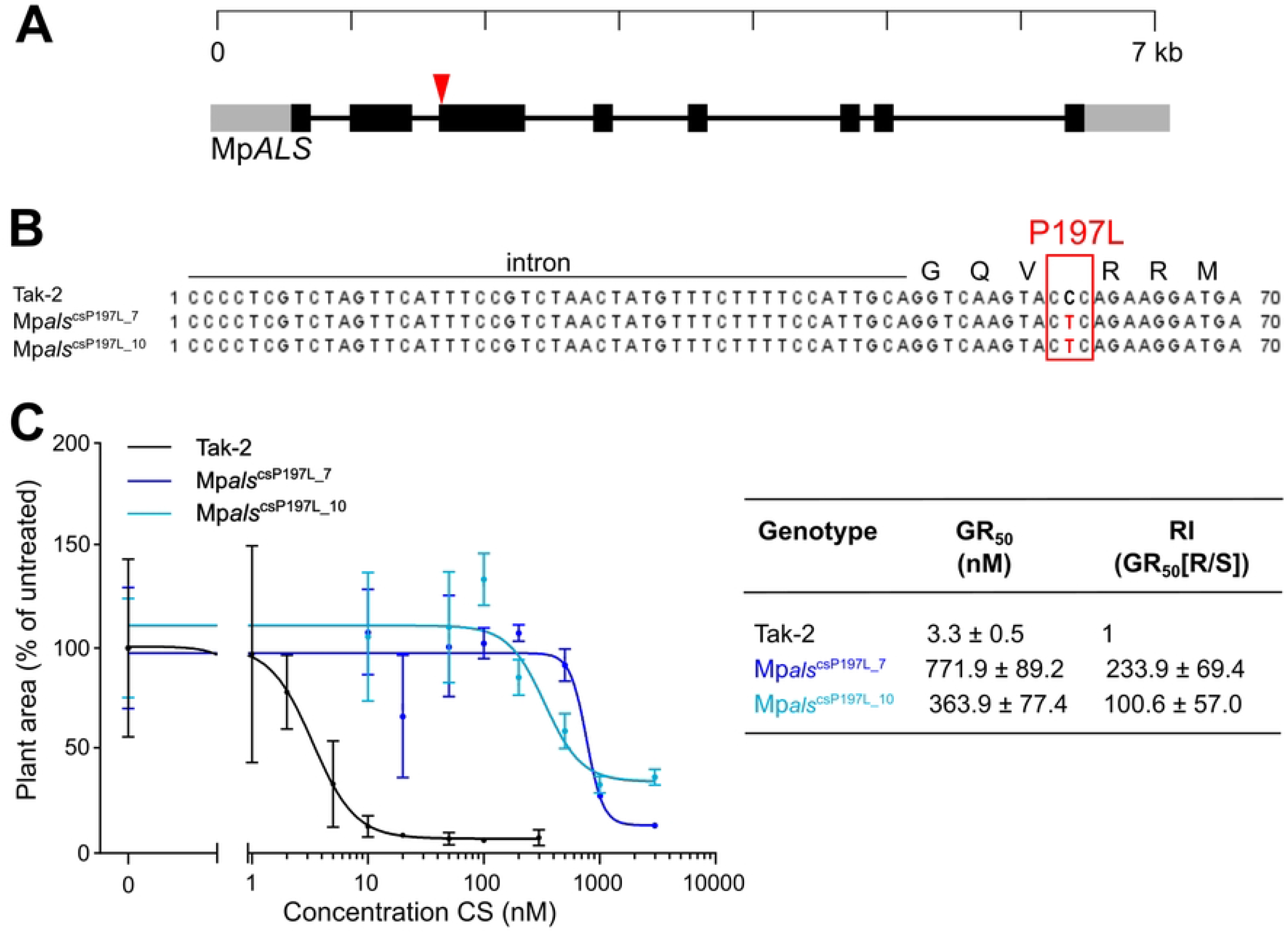
The P197L mutation in Mp*ALS* confers resistance to chlorsulfuron. **(A)** Diagram of the Mp*ALS* gene showing the location of the mutation in Mp*ALS* identified in lines Mp*als*^csP197L_7^ and Mp*als*^csP197L_10^ (arrow head). Untranslated regions (grey), exons (yellow). **(B)** Alignment of the sequenced region of Mp*ALS* showing the cysteine to thymine mutation which is predicted to cause an amino acid change of a proline to a leucine at amino acid position 197 of At*ALS* (position 207 in Mp*ALS*). **(C)** Growth assay of wild-type Tak-2, Mp*als*^csP197L_7^ and Mp*als*^csP197L_10^ lines grown on different concentrations of CS. Error bars are ± SD. N=5-10. GR_50_ is the herbicide concentration that causes a 50% reduction in growth. Resistance index (RI) = GR_50_ (Resistant line)/GR_50_ (Tak-2).

To quantify the degree of resistance of Mp*als*^*csP197_7L*^ and Mp*als*^*csP197L_10*^ mutant and wild-type plants were grown on media containing different concentrations of chlorsulfuron. The GR_50_ values of Mp*als*^*csP197L_7*^ and Mp*als*^*csP197L_10*^ were 771.9 nM (SD = 89.2) and 363.9 nM (SD = 77.4) respectively (Fig 2C). This was more than two orders of magnitude greater than wild-type which was 3.4 nM (SD = 1.0). While the lethal dose of chlorsulfuron for wild-type was 300 nM, we were unable to detect a lethal dose for the mutants in this experiment because mutant plants grew at the highest concentrations of chlorsulfuron applied (3 µM). Such high levels of resistance in mutants compared to wild-type is typical for mutations that alter the binding of the herbicide to its target (41,47,48).

The demonstration that mutations at the proline 197 codon confer resistance to chlorsulfuron is consistent with chlorsulfuron inhibiting ALS function in *M. polymorpha*.

### Mp*acetolactate synthase* (Mp*als*) complete loss of function mutants are lethal and are rescued by branched chain amino acid supplements

We demonstrate that the P197L mutation in Mp*ALS* confers resistance to chlorsulfuron and the lethality of chlorsulfuron in treated plants can be rescued with branched chain amino acids. This is consistent with ALS being the target of chlorsulfuron in *M. polymorpha*. Since ALS inhibition is lethal in *M. polymorpha*, we hypothesized that mutations in Mp*ALS* leading to a complete loss of ALS activity would not survive on standard media that lacked branched chain amino acids. We also predicted that Mp*als* loss of function mutants would be rescued by growing them in media with branched chain amino acids, in the same way that chlorsulfuron-lethality was suppressed by growing chlorsulfuron-treated wild-type plants in the presence of branched chain amino acids.

We generated Mp*als* mutants by CRISPR/Cas9 mutagenesis using two separate guide RNAs. The first guide RNA (F) was complementary to a sequence in the second exon, 5’ of the conserved thiamine pyrophosphate (TPP) binding domain. The second guide RNA (W) was complementary to a sequence near the beginning of the last exon. We predicted that mutations at these sites could lead to a total loss of function of the encoded protein. Plants were transformed with either the F or the W guide RNAs and plated on selective medium supplemented with branched chain amino acids. The medium was supplemented with branched chain amino acids because it was expected that at least some of the Mp*als* mutants would be auxotrophic for branched chain amino acids. 100 lines were selected from the transformation and the Mp*ALS* gene was sequenced in 31 plants. 16 of the sequenced transformants had mutations in Mp*ALS*. Four mutant lines – Mp*als*^*lofF12*^, Mp*als*^*lofF16*^, Mp*als*^*lofF32*^, Mp*als*^*lofF84*^ – with mutations in the regions of the gene complementary to the F guide RNA were identified. We predict that Mp*als*^*lofF16*^, Mp*als*^*lofF32*^, Mp*als*^*lofF84*^ are loss of function mutations because each has deletions that result in frame shifts in the coding sequence. We predict that Mp*als*^*lofF12*^ is either wild-type or a weak loss of function mutant because it harbours an in-frame insertion. Two mutant lines – Mp*als*^*lofW20*^ and Mp*als*^*lofW29*^ – were identified in the regions of the gene complementary to the W guide RNA. The mutations in the *MpALS* sequence in each of the six mutants are presented in Fig 3.

**Fig 3.**
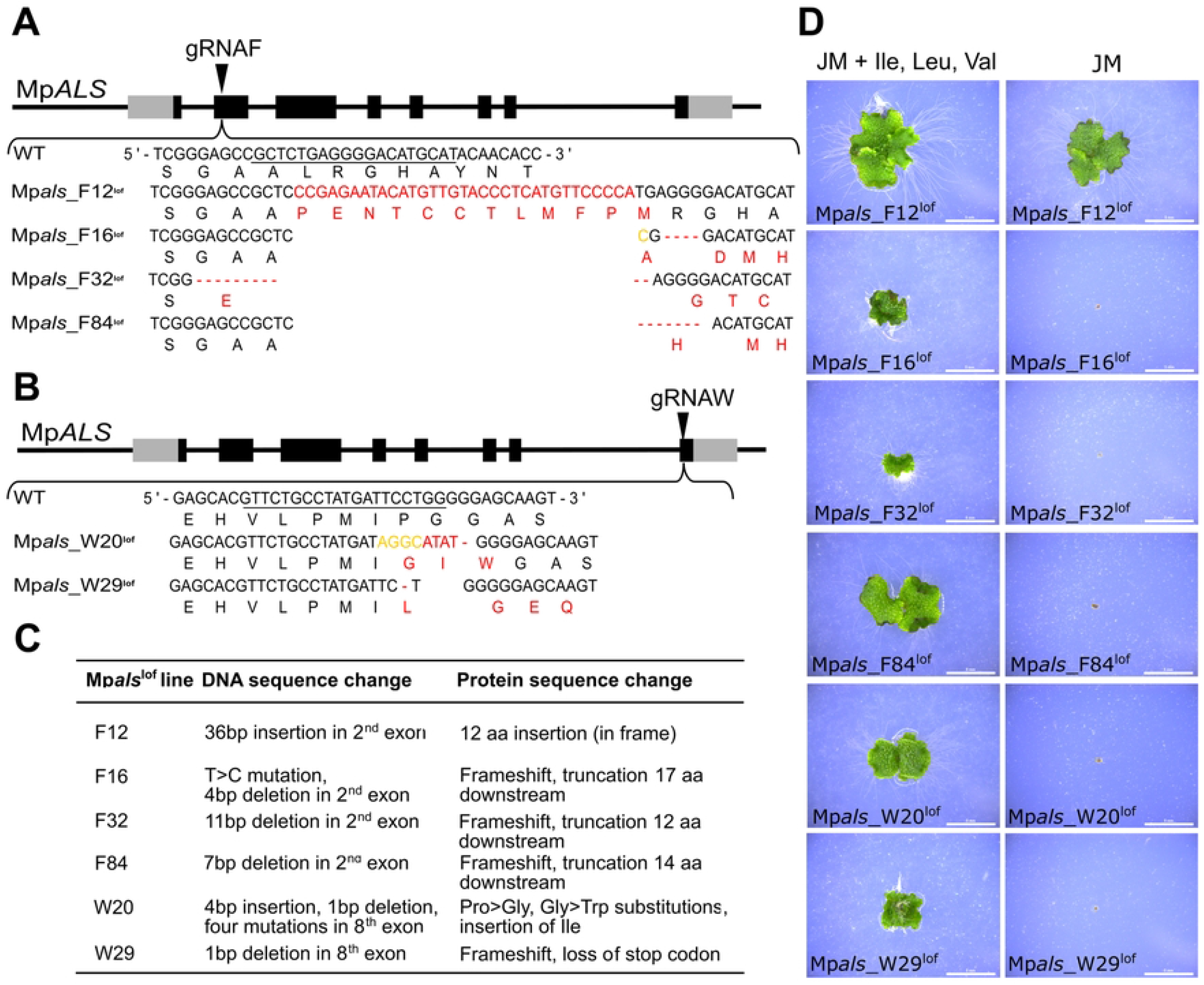
Six independent Mp*ALS* mutant lines were generated using CRISPR/Cas9 mediated targeted mutagenesis. **(A, B)** Mutations in Mp*ALS* were generated using a gRNA targeting the 2^nd^ exon (gRNAF) or the 8^th^ exon (gRNAW). The thiamine pyrophosphate binding domain is represented by grey boxes, all other exons are represented by black boxes. Mutations are indicated by yellow letters, insertions by red letters, and deletions by red dashes. **(C)** A description of the mutations generated by CRISPR/Cas9 and predicted amino acid changes in the 6 Mp*als*^*l*of^ mutants. **(D)** 18-day-old gemmalings grown on Johnsons’ medium with or without supplemented branched chain amino acids (800 µM). Scale bar = 5 mm.

Different Mp*als* mutants were grown on branched chain amino acids and gemmae from each line were harvested. These gemmae were then grown either in standard media (without branched chain amino acids) or in media supplemented with branched chain amino acids. The line containing the in-frame insertion – Mp*als*^*lof_F12*^ *–* grew in both the standard media and the branched chain amino acid-supplemented media (Fig 3D). This is consistent with the hypothesis that this mutant is prototrophic for branched chain amino acids as expected for a line with a wild-type Mp*ALS* or weak loss of function Mp*ALS* gene. By contrast, all five of the putative strong loss of function mutants grew on media containing branched chain amino acids, while none grew on standard media without the branched chain amino acid supplements. This suggests that these mutants are branched chain amino acid auxotrophs. These data demonstrate that loss of ALS function is lethal in *M. polymorpha* and the lethal phenotype can be suppressed by branched chain amino acid supplements. The observation that branched chain amino acid media supplementation suppresses the lethality of Mp*als* loss of function mutatations and chlorsulfuron treatment, is consistent with the hypothesis that chlorsulfuron inhibits ALS activity in *M. polymorpha*.

### Chlorsulfuron-treatment induces transcriptome changes and the accumulation of some non-branched chain amino acids

To determine if the presence of chlorsulfuron induced transcriptome changes we generated transcriptomes from plants that were grown on standard media for 8 days and were then transferred to chlorsulfuron-containing media for between 2 and 48 hours. Control plants were plants grown on standard media for 8 days and then transferred to standard media (with no chlorsulfuron) for 2-48 hours. Transcriptomes for each time point and treatment were generated from two technical replicates and each replicate consisted of six plants. There was a large difference in the transcriptomes of chlorsulfuron-treated and untreated controls. Steady state levels of 3093 mRNAs were changed by chlorsulfuron treatment. 1612 mRNAs were more abundant in at least one timepoint in chlorsulfuron-treated plants than in controls. Steady state levels of 1708 mRNAs were less abundant in at least one time point in the treated plants than in controls (Fig 4A). The differences in the transcriptomes of chlorsulfuron-treated and control plants are consistent with the hypothesis that chlorsulfuron-treatment induces transcriptome changes in *M. polymorpha*.

**Fig 4.**
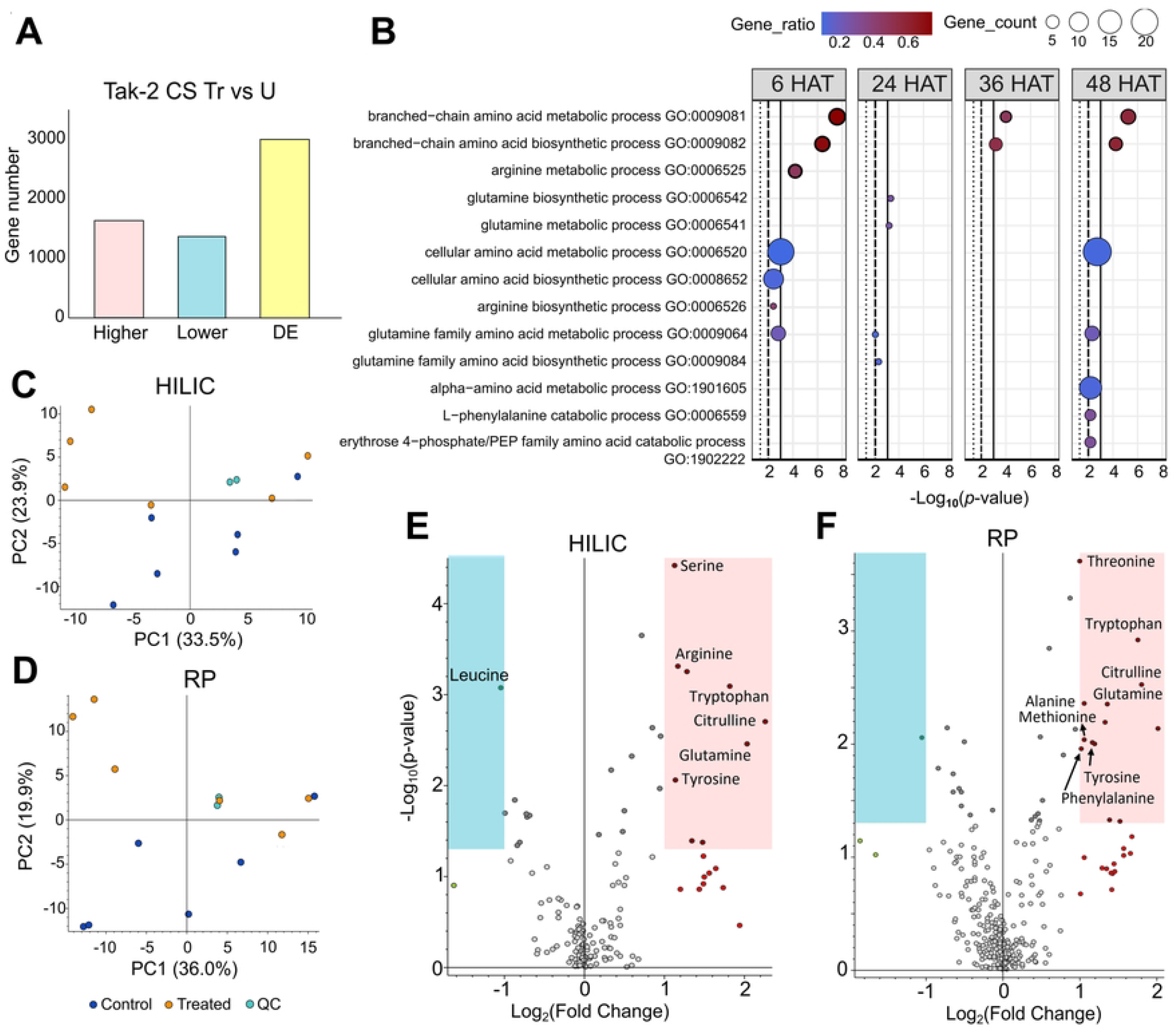
Chlorsulfuron treatment induces transcriptome changes and high levels of some non-branched chain amino acids. **(A)** Numbers of genes with altered mRNA steady state levels 0-48 h after chlorsulfuron treatment compared to control treatments. **(B)** Enriched Gene Ontology (GO) terms of mRNAs that were more abundant in chlorsulfuron-treated plants compared to untreated plants. Cut-off lines drawn at equivalents of p=0.05, p=0.01, p=0.001. **(C-E)** Non-targeted metabolomic analysis of wild-type plants treated for 24 h with 100 nM chlorsulfuron and untreated plants. Samples were analysed by LC-MS/MS using hydrophilic interaction liquid chromatography (HILIC) and reversed phase chromatography (RP). Principal component analyses of control (C1-6) and herbicide-treated (H1-6) samples using HILIC **(C)** and RP **(D)**. Volcano plots of differentially abundant metabolites in chlorsulfuron-treated plants compared to untreated plants using HILIC **(E)** and RP **(F)**. Blue box indicates significantly lower levels of metabolites and red box indicates significantly higher levels of metabolites. Significance based on one-way ANOVA followed by Tukey HSD post-hoc test.

To identify physiological processes that were impacted by the chlorsulfuron-induced transcriptome changes we carried out an analysis of Gene Ontology (GO) terms that were enriched among the 1612 mRNAs with higher steady state levels in chlorsulfuron-treated plants compared to controls. This showed that GO terms associated with branched chain amino acid biosynthesis and branched chain amino acid metabolism are significantly enriched at 6 hours (*p* < 1.10^−6^ and *p* < 1.10^−7^), 36 hours (*p* < 0.001 and *p* < 1.10^−5^) and 48 hours (*p* < 1.10^−4^ and *p* < 1.10^−5^) after chlorsulfuron-treatment compared to other GO terms (Fig 4B). This is consistent with the hypothesis that chlorsulfuron treatment blocks branched chain amino acid synthesis in *M. polymorpha* because inhibition of the pathway results in an increase in the mRNA levels of genes that code for proteins involved in branched chain amino acid synthesis.

The GO term enrichment analysis also demonstrates that GO terms for amino acid biosynthesis and metabolism are enriched after 6 hours chlorsulfuron treatment in mRNAs that are more abundant in chlorsulfuron-treated plants than controls (Fig 4B). These transcriptome profiles predict that levels of some non-branched chain amino acids would be higher in chlorsulfuron-treated plants than in untreated controls. To test this hypothesis, we measured the levels of amino acids (and other metabolites) in chlorsulfuron-treated and untreated controls. Plants were grown for 11 days and treated with 100 nM chlorsulfuron for 24 hours before being extracted. Control plants were transferred to standard media for 24 hours before extraction. Using two independent separation methods – hydrophilic interaction liquid chromatography and reversed phase chromatography – we found that levels of threonine, tryptophan, arginine, serine, alanine, tyrosine, glutamine, citrulline and phenylaniline are higher in chlorsulfuron-treated plants than in the untreated controls (Fig 4 C-F). This is consistent with the transcriptome profiles observed in chlorsulfuron-treated plants. This suggests that inhibition of ALS causes changes not only in the branched chain amino acid pathway but in other amino acid pathways and therefore may have wide ranging physiological impacts on the plant.

We conclude that inhibition of branched chain amino acid synthesis by chlorsulfuron results in an increase in the steady state levels of mRNAs for enzymes involved in amino acid synthesis, possibly due to a branched chain amino acid deficiency-induced transcriptional response. These transcriptome changes may have led to increased levels of amino acids such as threonine, tryptophan, arginine, serine, alanine, tyrosine, glutamine, and phenylaniline.

### Induction of gene expression by chlorsulfuron requires the inhibition of the herbicide-sensitive target acetolactate synthase

It has been hypothesized that a sensing mechanism initiates the detoxification of herbicides in plants (26). According to this hypothesis, a sensing mechanism detects the herbicide, and the sensor programs transcriptome changes that increase the expression of genes encoding proteins that chemically modify herbicides (detoxification). An alternative hypothesis is that there is no specific herbicide sensing mechanism and therefore no inducible system of herbicide resistance. If the difference between herbicide-treated and untreated transcriptomes depends on a sensing mechanism, the differences should also be observed in chlorsulfuron-treated plants even if chlorsulfuron cannot inhibit its target, as in Mp*als*^*cs*^ target site resistant mutants. If, on the other hand, the transcriptome changes are the result of metabolic changes caused by inhibition of ALS activity, then the transcriptome changes should not be observed in Mp*als*^*cs*^ target site resistant mutants.

To determine if the chlorsulfuron-induced transcriptome changes observed in wild-type were also observed in chlorsulfuron target site resistant mutants, we generated transcriptomes from chlorsulfuron-treated and untreated Mp*als*^*csP197L_7*^ and Mp*als*^*csP197L_10*^ target site resistant mutants (Fig 5). While chlorsulfuron-treatment of wild-type plants resulted in large changes in the steady state levels of over 3093 mRNAs, the steady state levels of most mRNAs were the same in chlorsulfuron-treated and untreated Mp*als*^*cs*^ mutants (Fig 5B). 21 mRNAs were more abundant and 16 less abundant in chlorsulfuron-treated Mp*als*^*csP197L_7*^ plants compared to untreated controls. Similarly, 2 mRNAs were more abundant and 1 less abundant in chlorsulfuron-treated Mp*als*^*csP197L_10*^ plants compared to untreated controls (Fig 5B). Importantly, no single mRNA changed abundance in every background – wild-type, Mp*als*^*csP197L_7*^ and Mp*als*^*csP197L_10*^ – when treated with chlorsulfuron compared to untreated controls (Fig 5C). These data indicate that the chlorsulfuron-induced transcriptome change observed in wild-type is not observed in Mp*als*^*csP197L_7*^ or Mp*als*^*csP197L_10*^ mutants.

**Fig 5.**
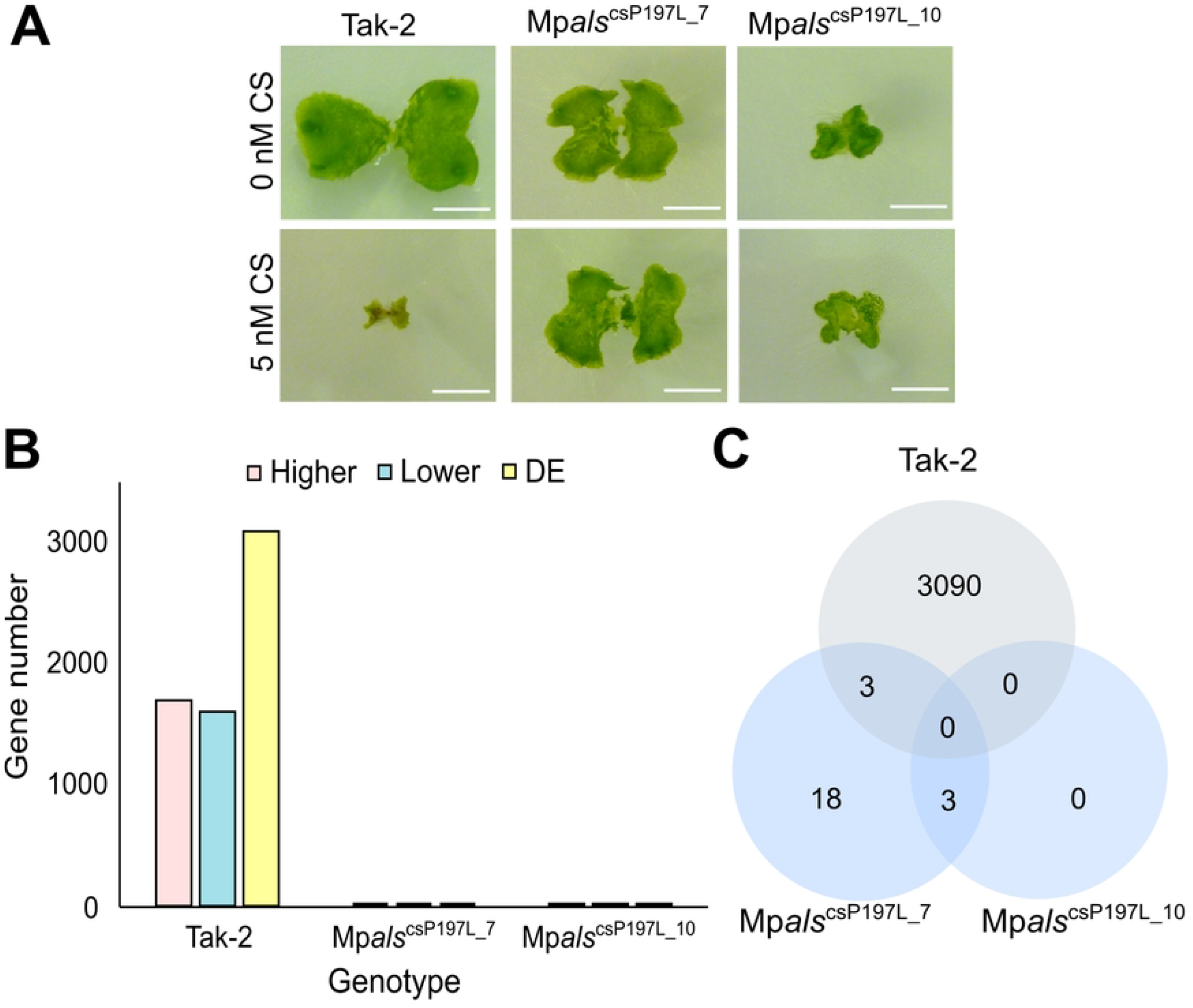
Chlorsulfuron treatment does not alter gene mRNA steady state levels in Mp*als*^chlrP197L^ mutants. **(A)** Wild-type, Mp*als*^*chlrP197_7*^ and Mp*als*^*chlrP197_10*^ lines grown for 7 days on untreated media then transferred to media containing 0 or 5 nM CS for 7 days. Scale bar = 5 mm. **(B)** Differentially expressed genes (DEGs) between CS-treated and untreated plants in each genotype. Blue bar represents genes with higher mRNA steady state levels, green bar represents genes with lower mRNA steady state levels, and yellow bar represents genes with either higher or lower mRNA steady state levels in CS-treated plants compared to untreated plants. **(C)** Number of differentially expressed genes in CS treated plants compared to untreated plants between Tak-2, Mp*als*^*chlrP197_7*^ and Mp*als*^*chlrP197_10*^.

These data are not consistent with chlorsulfuron itself being sensed by the plant and this sensing mechanism initiating a transcriptome response. Instead, the data are consistent with the transcriptome changes induced by chlorsulfuron treatment being the result of the inactivation of the herbicide target, ALS. The change in gene expression caused by chlorsulfuron-treatment is likely a result of the metabolic changes caused, directly and indirectly, by the inhibition of ALS by the herbicide.

### Ectopic overexpression of genes encoding cytochrome P450 monooxygenases and glutathione S-transferases does not confer resistance to chlorsulfuron

If the chlorsulfuron-induced transcriptome changes result from the inhibition of ALS and its physiological consequences – and not from a chlorsulfuron-sensing mechanism that induces transcriptome changes that lead to the production of enzymes that detoxify chlorsulfuron – we predicted that genes encoding enzymes induced by exposure to chlorsulfuron would not form part of such a sensing mechanism and would therefore not detoxify the herbicide.

To test this hypothesis, we ectopically overexpressed four genes whose steady state levels of mRNA abundance increased upon chlorsulfuron treatment (Table 1). To maximize the chances of identifying a gene with detoxifying potential, we selected members of the cytochrome P450 monooxygenases and glutathione transferases, because overexpression of members of these families in mutant weeds confers herbicide resistance (49). To further increase the chances of identifying members of the GST and CYP families that might detoxify chlorsulfuron, we selected genes from GST and CYP clades that had previously been shown to increase herbicide tolerance when overexpressed in herbicide resistant weeds (49–52). Using these criteria, we selected Mp*CYP813A5* and Mp*CYP822A1* from the cytochrome P450 clan 72 and clan 71 respectively, and Mp*GSTF15* and Mp*GSTF10* from the glutathione transferase ϕ clade.

**Table 1.**
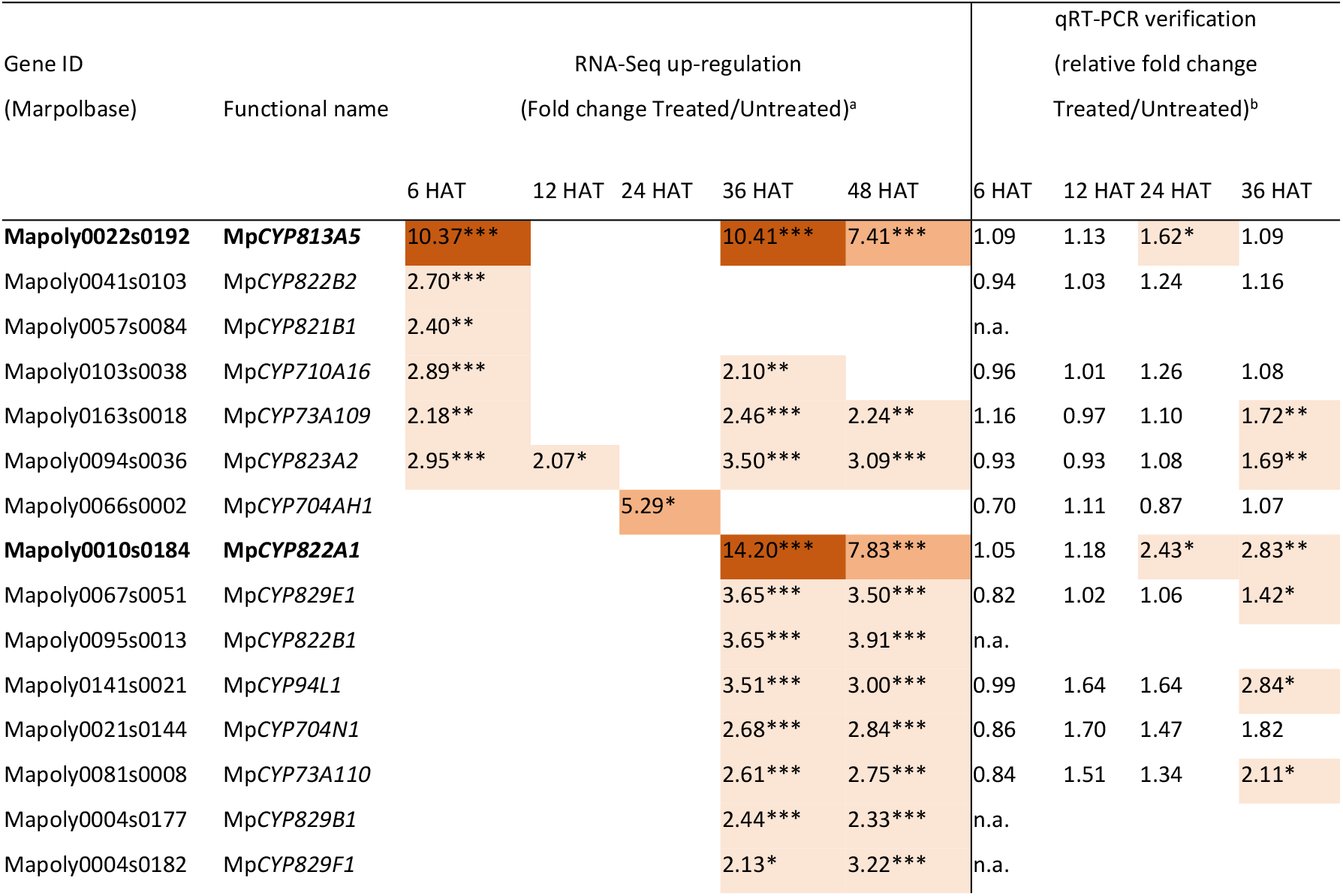
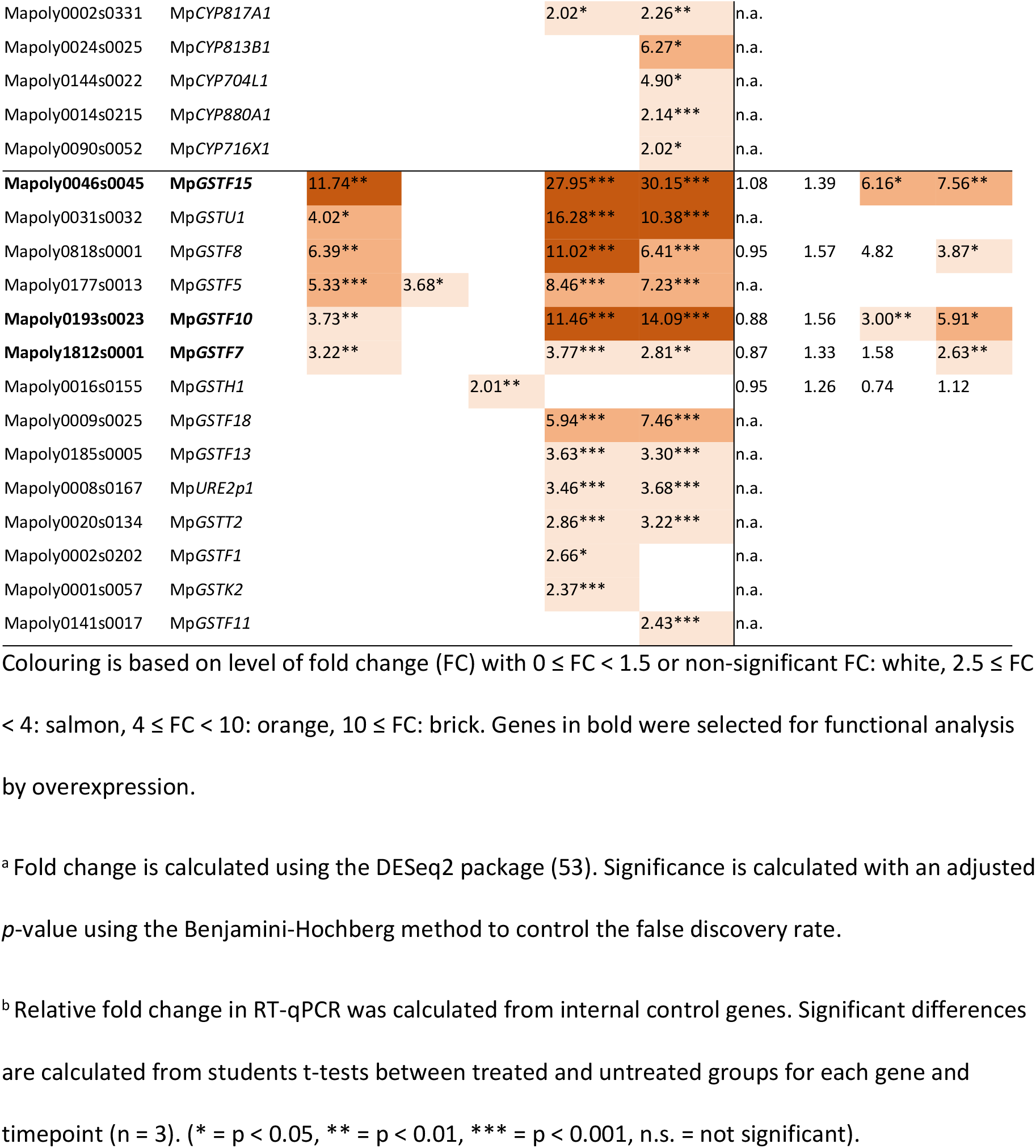
Several cytochrome P450 monooxygenases and glutathione S-transferases had higher mRNA steady state levels in CS-treated compared to untreated plants in both RNA-Seq and qRT-PCR experiments.

Wild-type spores were transformed with overexpression vectors containing each gene under the transcriptional control of the Mp*ELONGATIONFACTOR1α* promoter (*proEF1α*) using the *NOS* terminator. Controls were generated by transforming wild-type spores with the empty vector (construct without the CYP or GST coding sequence). Transformed plants were grown, RNA isolated and the steady state levels of the respective CYP and GST mRNA measured using quantitative reverse transcription polymerase chain reaction (RT-qPCR). Steady state levels of Mp*CYP813A5* and Mp*CYP822A1* mRNA were 4000 times higher and 60 times higher in *proEF1α*:Mp*CYP813A5*_7 and *proEF1α*:Mp*CYP822A1*_3 transformed plants than in empty vector controls respectively (Fig 6 A). Steady state levels of MpGSTF15 and MpGSTF10 from mRNA were 2.5 times higher and 300 times higher in *proEF1α*:Mp*GSTF15*_5 and *proEF1α*:Mp*GSTF10*_17 transformed plants than in empty vector controls respectively (Fig 6 A).

**Fig 6.**
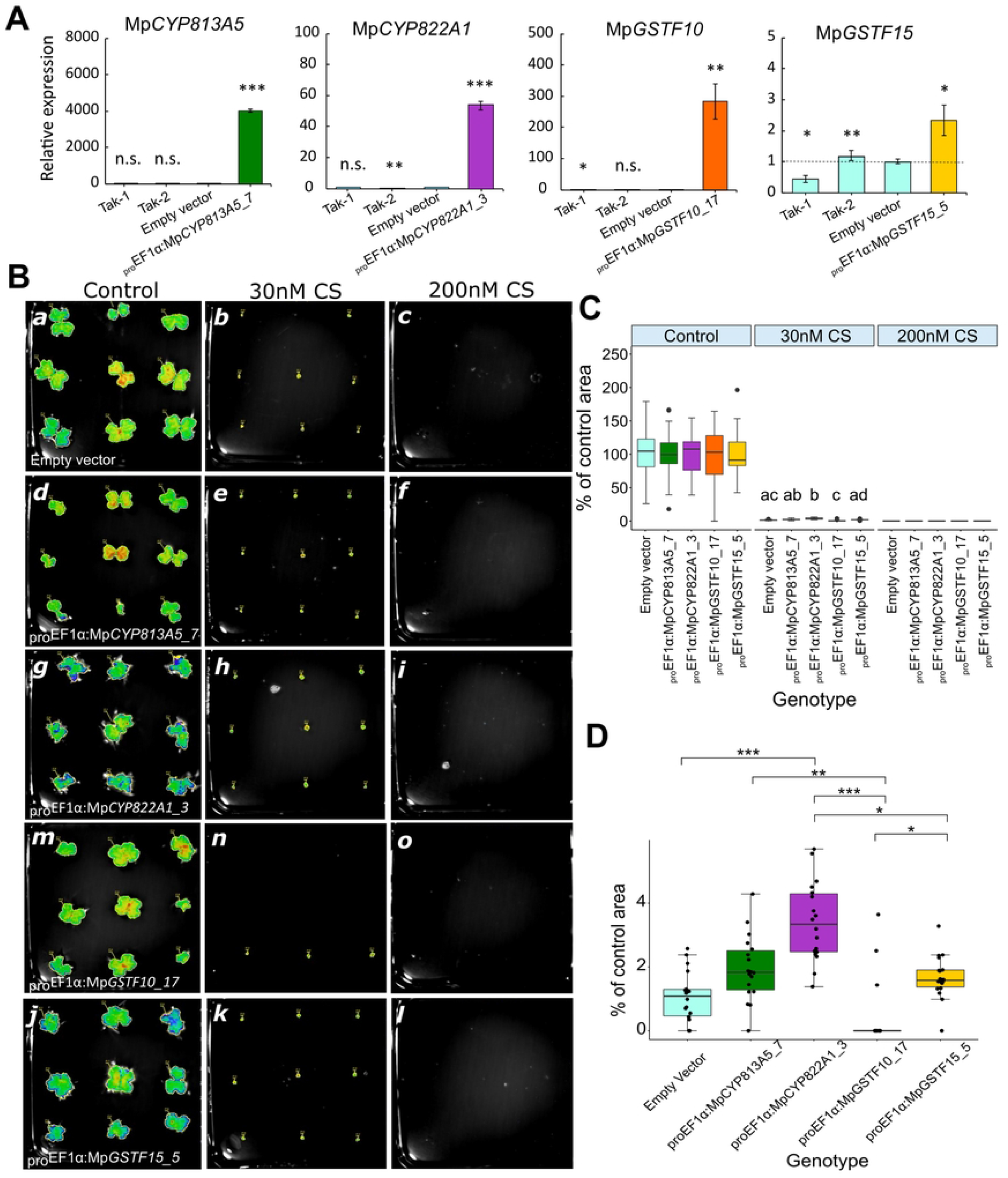
Ectopic overexpression of genes encoding cytochrome P450 monooxygenases and glutathione S-transferases does not confer resistance to chlorsulfuron. **(A)** Relative mRNA abundance in 12-day-old gemmalings of Mp*CYP813A5*, Mp*CYP822A1*, Mp*GSTF10* and Mp*GSTF15* over-expressing lines and wild-type (Tak-1 and Tak-2) was compared to relative mRNA abundance in empty vector controls (dotted line, n = 3). mRNA abundance is given as x-fold change. Statistical significance based on student t-test is given for each line compared to empty vector control. *=p < 0.05, ** = p < 0.01, *** = p < 0.001, n.s. = not significant. Error bars are ± SD. **(B)** 14-day-old gemmalings of an empty vector line (B, a-c) and of Mp*CYP813A5*, Mp*CYP822A1*, Mp*GSTF10* and Mp*GSTF15* over-expressing lines (B, d-l) grown on untreated solid medium (Control), 30 nM CS and 200 nM CS. Images taken on a Berthold NightOwl. Chlorophyll autofluorescence is used to measure gemmaling area (mm^2^) which is treated as a proxy for growth. Images of whole 120×120 mm square petri dish. **(C)** Box and whisker plot where whiskers are first and fourth quartiles, boxes represent second and third quartiles, and the horizontal line is the median. N = 18. Statistical significance is measured by a Kruskal-Wallis test. Significant groups determined by a Nemenyi post-hoc test. * = p<0.05, ** = p < 0.01, *** = p < 0.001, n.s. = not significant. Error bars are ± SD. The same data for the 30 nM CS treatment is shown on a smaller y axis in **(D)**.

Gemmae from control and transformed plants were grown on 0 mM (control), 30 nM and 200 nM chlorsulfuron. The growth of all transformed genotypes was reduced slightly compared to the empty vector control when grown on media without chlorsulfuron (Fig 6 B, C). Therefore, to control for within-genotype growth differences, plant area was presented as percentage of area of untreated plants for each genotype (Fig 6 D). No genotypes grew on media containing 200 mM chlorsulfuron, indicating that the plants transformed with the CYP and GST overexpressing constructs were sensitive to the herbicide (Fig 6 B, C). There was almost no growth of any genotype on 30 mM, but *proEF1α*:Mp*CYP822A1*_3 transformed plants grew slightly more than the control (*p* < 0.001) (Fig 6B, C, D). The *proEF1α*:Mp*CYP822A1*_3 transformed plants grew to 3.4% (SD = 1.2) of the area of untreated plants, while the empty vector grew to 1.1% (SD = 0.8) area of untreated plants. However, this percentage growth is very small compared to each genotype grown in the absence of chlorsulfuron, therefore Mp*CYP822A1* overexpression does not restore growth of chlorsulfuron-treated plants to untreated levels

We conclude that ectopic overexpression of two CYP-encoding genes and two GST-encoding genes did not confer chlorsulfuron tolerance. These data are consistent with the hypothesis that genes whose steady state mRNA levels increase on chlorsulfuron-treatment are not involved in herbicide detoxification.

## Discussion

We show that the herbicide chlorsulfuron induces a transcriptome response in sensitive *M. polymorpha* plants. However, this transcriptome response does not occur in target-site-resistant mutants (*Mpals*^*cs*^) treated with chlorsulfuron. The observation that the transcriptome change does not occur in mutants where the chlorsulfuron cannot inhibit the activity of its target, the ALS enzyme, suggests that inhibition of ALS is required for the transcriptome response. It also suggests that the presence of chlorsulfuron alone is not sufficient to induce the transcriptome response. If the presence of chlorsulfuron is insufficient to induce a transcriptome response, it suggests that the herbicide molecule is not sensed by the plant. We conclude that a mechanism that senses chlorsulfuron does not exist in *M. polymorpha*, and the observed transcriptome response is the result of direct and or indirect effects of the inhibition of ALS by chlorsulfuron. If this observation is holds true for other plant species and with other herbicides, it suggests that a herbicide-sensing system that induces herbicide detoxification mechanisms does not exist in plants. We hypothesize that a herbicide-sensing system that induces transcriptional changes that promote the production of proteins active in herbicide detoxification does not exist in *M. polymorpha*, nor perhaps in other plants.

There is abundant evidence in the literature that treatment of plants with herbicides results in changes in steady state mRNA levels that are consistent with our data. However, the underlying cause of these herbicide-induced transcriptome changes has remained unclear. The transcriptome changes could have been due to the existence of a sensing mechanism that perceives the presence of a herbicide, which then activates a signalling pathway resulting in changes in gene expression as observed in the transcriptomes. Alternatively, the transcriptome changes could also have been the result of the inhibition of the target protein by the herbicide. In the research reported here, we have been able to distinguish between these possibilities. The data demonstrate that the chlorsulfuron-induced changes only occur if the herbicide target, the ALS enzyme, is sensitive to the herbicide. If the herbicide target is insensitive to the enzyme, the transcriptome change does not occur. Our data are therefore consistent with transcriptome data reported in the literature, but go a step further to show that the transcriptome change depends on the defective function of the herbicide target.

While we report that a chlorsulfuron sensing system does not exist in *M. polymorpha* and we hypothesize that herbicide sensing mechanisms do not exist in plants, since it has been shown that transcriptome responses to ALS inhibitors are highly conserved across plant species (54), it is formally possible that such a sensing mechanism may exist in some plants for some herbicides. If so, we expect the presence of herbicides would activate a signalling pathway that would cause transcriptome changes and the production or activation of proteins that chemically modify the herbicide and reduce its toxicity. Such a herbicide-plant interaction remains to be demonstrated.

Our conclusion – that a herbicide sensing mechanism that induces herbicide detoxification does not exist – is consistent with observations from the field. Resistance to herbicides has evolved many times in the field. Non target site resistance (NTSR) is a form of herbicide tolerance which prevents the active herbicide from coming into contact with its target, either through the chemical modification of the compound or the sequestration of the compound so that it no longer has access to the herbicide target, and therefore cannot inhibit its biological function. NTSR is often the result of gene overexpression. These genes may encode proteins that oxidise the herbicide (e.g. cytochrome P450 monooxygenases), conjugate the herbicide to glutathione (e.g. glutathione transferases) or to saccharides (e.g. glycosl-transferases) or transport the herbicide across a membrane (e.g. ABC transporters). Their overexpression is the result of a mutation that increases their expression levels compared to “wild-type”, herbicide sensitive weeds. These overexpressing, mutant alleles, would arise rarely in the field. However, these rare mutations would be selected for in the presence of intense herbicide selection. They could become dominant alleles in populations undergoing prolonged herbicide selection. While these resistance-conferring alleles contribute to resistance in the field, our data suggest that they are not part of an inducible mechanism that detoxifies herbicides.

## General Conclusion

These data indicate that while chlorsulfuron-treatment results in transcriptome changes, these changes are not the result of a herbicide-sensing mechanism. Instead, the transcriptome changes depend on the inactivation of the chlorsulfuron target. This is consistent with the hypothesis that transcriptome changes result from the disrupted physiology caused by the inhibition of ALS activity and are not specifically related to herbicide detoxification.

## Materials and methods

### Plant materials and growth conditions

*M. polymorpha* accessions Takaragaike-1 (Tak-1, male) and Takaragaike-2 (Tak-2, female) were used as wild-type plants (55). Mp*als*^*csP197L*^, Mp*als*^lof^, Mp*CYP813A5*^*GOF*^, Mp*CYP822A1*^*GOF*^, Mp*GSTF10*^GOF^ and Mp*GSTF15*^GOF^ lines were generated as described below. Plants were propagated asexually from gemmae and grown in light cabinets under 30 μmolm^−2^s^−1^ continuous white light at 23 °C.

Plants used in the metabolomic analyses, transcriptome analysis, quantitative real time PCR verification of transcriptome data and chlorsulfuron dose response assay were grown on 100×21 mm round petri dishes on top of autoclaved cellulose discs (A.A. Packaging Ltd, Preston, UK) placed on the surface of 30 ml Johnson’s medium (56) with 0.8% agar. Plants were transferred from the herbicide-free medium to herbicide-containing medium by lifting the discs on which plants were growing from one plate to another. Plants used in the other experiments were grown on 120×120 mm square petri dishes with 60 mL half strength Gamborg medium (57) with 1% sucrose and 0.8% agar.

To generate spores for transformations, Tak-1 and Tak-2 plants were grown on a 1:3 mixture of vermiculite and John Innes No.2 compost and incubated in a growth chamber under a 16 h light:8 h dark photoperiod at 23°C. Gametophore production was induced under far red light irradiation (58).

### Generation of Mp*als*^*csP197L*^ lines

Mp*als*^*csP197L*^ mutant *M. polymorpha* plants were generated via UV-B mutagenesis. 10^6^ spores were mutated under a UV-B wavelength of 302 nm and grown on 140 nM (0.05 ppm) chlorsulfuron under continuous light. Resistant plants were selected after 14 days.

### Sequencing Mp*ALS* in wild-type and Mp*als*^*csP197L*^ plants

To determine the mutations in the ALS gene of Mp*als*^*csP197L*^ lines, whole plant DNA was extracted from two mutants (Mp*als*^*csP197L_7*^ and Mp*als*^*csP197L_10*^) and wild-type (Tak-2) using the CTAB method adapted from (59). DNA concentration was measured on a Nanodrop ND-1000 spectrophotometer.

A 140 bp region of the *M. polymorpha ALS* gene that contains the most commonly found mutations conferring resistance to ALS-inhibitors was amplified by polymerase chain reaction (PCR) using the PCRBIO HiFi polymerase and buffer (PCR Biosystems, Cat. No. PB10.41-02). The amplification was carried out using a BIO-RAD Dyad Peltier thermal cycler. Each reaction was conducted in a 50 μl mixture, consisting of 10 μl reaction buffer, 2 μl of forward primer, 2 μl of reverse primer, 5 μl of DNA, 0.5 μl of HiFi Polymerase, and 30.5 μl of DNase free milli-Q H2O, with three replicates per sample. The PCR program consisted of 1 min initial denaturation phase at 95 °C, followed by 32 cycles, with each cycle consisting of a 15 s incubation time at 95 °C, a 15 s incubation at 56 °C annealing temperature, and a 30s extension time at 72 °C. Amplified DNA was extracted from the gel using the QIAquick Gel Extraction Kit (Qiagen, Hilden, Germany). The quantity of extracted DNA was measured on a Nanodrop ND-1000 spectrophotometer. Samples were sequenced by Sanger sequencing (Source Bioscience, Oxford, UK). The primers used for PCR amplification and sequencing are listed below.

### Chlorsulfuron effect on *M. polymorpha* growth

A whole plant dose-response experiment was carried out to determine the sensitivity of wild-type and target-site resistant plants to chlorsulfuron. Chlorsulfuron, 1-(2-chlorophenylsulfonyl)-3-(4-methyoxy-6-methyl-1,3,5-triazin-2-yl)urea, was supplied by Sigma Aldrich and diluted in milli-Q H2O, then filter sterilized with a 0.2 µM sterile syringe filter. Chlorsulfuron was pipetted into melted medium and mixed thoroughly before pouring into plates. The female *M. polymorpha* accession (Tak-2) was chosen as the susceptible biotype to chlorsulfuron because it tends to grow flatter on the medium than Tak-1 making it easier to image. Tak-2, Mp*als*^*csP197L_7*^ and Mp*als*^*csP197L_10*^ gemmae were grown for 7 days then transferred onto herbicide treated plates and replaced in the growth cabinet for 7 days before imaging. Tak-2 plants were treated with ten herbicide doses: 0 nM, 0.1 nM, 0.2 nM, 0.5 nM, 1 nM, 1.5 nM, 2.5 nM, 5 nM, 10 nM, and 30 nM chlorsulfuron. Mp*als*^*csP197L*^ plants were treated with ten herbicide doses: 1 nM, 2 nM, 5 nM, 10 nM, 20 nM, 50 nM, 100 nM and 300 nM chlorsulfuron. There were two replicate plates per herbicide dose and five plants grown per plate. The plants were imaged using a Panasonic Lumix camera (model no. DMC-FS7) on day 7 after transfer to herbicide plates. Lateral growth was quantified using Image J. Data were expressed as % of the average area of untreated plants and were plotted with a non-linear fit, sigmoidal, four-parameter regression curve using GraphPad Prism 9.1.0 (GraphPad Software, Inc., San Diego, CA, USA). The herbicide dose resulting in 50% growth reduction (GR_50_) was calculated as follows:

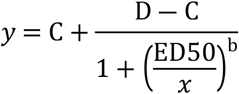

In which C is the lower limit, D is the upper limit, ED_50_ is the effective dose which reduced growth by 50%, and b is the slope at ED_50_. Resistance index was then calculated as follows: 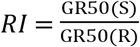

### Branched chain amino acid growth assay

Tak-2 gemmae were grown in petri dishes on solid medium supplemented with 800 µM branched chain amino acids, 800 µM branched chain amino acids and 100 nM CS, with 100nM CS only or on unsupplemented medium and grown for 14 days. Whole plates were imaged using a Berthold NightOwl II LB 983 *In Vivo* imaging system (Berthold, Bad, Wildbad, Germany) which detects chlorophyll autofluorescence (560 nm). Images were taken after exposing the plants to 120 s white light. The area (mm^2^) of living tissue was measured using indiGo™ software (Berthold, Bad Wildbad, Germany). Statistical significance was based on Kruskal-Wallis test, with groups determined by Nemenyi post-hoc test.

### Sample extraction for metabolomic analysis

11-day old Tak-2 plants were transferred to medium containing 100 nM CS or untreated medium for 24 hours, then extracted. Approximately 150 mg of fresh plant tissue per sample was collected and frozen immediately in liquid N2. 6 biological replicates were collected per treatment group. Metabolite extraction was as follows. 500 µL methanol:acetonitrile:H_2_O (2:2:1, v/v) (stored at −20 °C) was added to each sample. Plant material was then homogenised for 2 min in a tissuelyser (Qiagen) using adaptors kept at −70 °C. Samples were incubated for 1 h at −20 °C followed by centrifugation at 14,000 rpm at 4 °C. The supernatant was removed and placed in a new Eppendorf tube, and stored at −20 °C. 400 <u>µ</u>L of 80% methanol:water (stored at −20 °C) was added to the precipitate then vortexed for 60 s at 4-8 °C. Samples were incubated for 1 h at −20 °C then centrifuged for 14,000 rpm at 4 °C. The supernatant from the first extraction was combined with the second extraction, and the samples were stored for 2 h at −20 °C. Samples were centrifuged one final time at 14,000 rpm for 10 min at 4 °C. The supernatant was transferred to a new Eppendorf tube, frozen in liquid N2 then stored at −80 °C.

### Targeted metabolomic analysis

Liquid chromatography-tandem mass spectrometry (LC-MS/MS) analysis of samples from chlorsulfuron-treated and untreated plants was performed by the Metabolomics Facility at Vienna BioCenter Core Facilities (VBCF), member of the Vienna BioCenter (VBC), Austria and funded by the City of Vienna through the Vienna Business Agency. For the analysis of branched chain amino acids and their precursors, 1 μL of the metabolite extract was injected on a Kinetex C8 column (100 Å, 100 × 2.1 mm), employing a flow rate of 80 μL/min. A 5-min-long gradient from 95% A (0.1 % formic acid in water) to 60% B (0.1 % formic acid in acetonitrile) was used for the separation. The HPLC (RSLC ultimate 3000; Thermo Fisher Scientific) was directly coupled to a TSQ Altis mass spectrometer (Thermo Fisher Scientific) via electrospray ionization. The following SRM (selected reaction monitoring) transitions were used in the positive ion mode: m/z 118.1 to m/z 72 (valine), m/z 132.1 to m/z 86.1 (isoleucine and leucine), m/z 205.1 to m/z 188 (tryptophan). Isopropylmalate wash was measured in the negative ion mode using the transitions m/z 175.1 to m/z 115 and m/z 118.1 to m/z 113. Optimal collision energies and retention times were determined and validated for each metabolite with the respective authentic standard. Other amino acid precursor metabolites (e.g. 2-ketobutyric acid) were targeted using the respective SRMs but only detected in the mixture of authentic standards. The data was manually interpreted using Trace Finder (Thermo Fisher Scientific).

### Non-targeted metabolomic analysis

For the non-targeted detection of metabolites in chlorsulfuron treated and untreated plants, samples were analysed by LC-MS/MS using the hydrophilic interaction liquid chromatography (HILIC) the reversed phase chromatography (RP) separation methods. 1 μL of each sample was injected independently onto two different phase systems, on a SeQuant ZIC-pHILIC HPLC column (Merck, 100 × 2.1 mm; 5 μm) and on a C18-column (Waters, ACQUITY UPLC HSS T3 150 × 2.1; 1.8 μm). 6 µL of each sample was pooled into a quality control (QC) sample used for normalization. Separation was performed with a flow rate of 100 μl/min employing an Ultimate 3000 HPLC system (Thermo Fisher Scientific, Germany). In HILIC (hydrophilic interaction liquid chromatography) a 25 min gradient from 10% to 80% B was used (A: acetonitrile (ACN); B: 25 mM ammonium bicarbonate in water) and in reversed phase chromatography (RP) a gradient from 1% to 90% B in (A: 0.1% formic acid (FA) in water; B: 0.1% FA in ACN). The HPLC was coupled via electrospray ionization to the mass spectrometer. Metabolites were ionized via electrospray ionization in polarity switching mode, acquiring high-resolution tandem mass spectrometry data on a Q-Exactive Focus (Thermo Fisher Scientific, Germany) in data-dependent acquisition mode. Data sets were processed by Compound Discoverer (Thermo Fisher Scientific), searching our in-house library and publicly available spectral libraries with a mass accuracy of 3 ppm for precursor masses and 10 ppm for fragment ion masses.

### gRNA design and vector construction for the generation of Mp*als* loss of function alleles

The generation of mutations in the Mp*ALS* gene using the CRISPR/Cas9 system followed the method developed by (60). Two short guide RNAs (gRNAs) gRNA_W (GTTCTGCCTATGATTCCTGG, complementary to nucleotides 1962 to 1984 of the coding sequence) and gRNA_F (CTCTGAGGGGACATGCATAC, complementary to the reverse strand of nucleotides 133 to 153 of the coding sequence) were designed to target two different locations in the Mp*ALS* DNA sequence. gRNAs must be upstream (5’) of a PAM sequence (NGG) and contain a protospacer or ‘target’ sequence consisting of 20 nucleotides that are unique to Mp*ALS*. gRNAs were designed using the CRISPR-P website (61). Individual gRNAs were cloned in the MpU6-1pro:gRNA expression vector using BsaI overhangs. MpU6-1pro:gRNA was combined with the Cas9 expression vector, MpGE010 via LR recombination. Vectors were transformed into *Escherichia coli* One Shot OmniMAX 2 T1 (ThermoFisher Cat# C854003). CRISPR-Cas9 expression vectors carrying gRNA_W or gRNA_F were transformed in *Agrobacterium tumefaciens* (GV3101). *Agrobacterium*-mediated spore transformation followed previously described methods (55), with the additional step of supplementing 800 μM isoleucine, leucine and valine to the *Agrobacterium* and spore liquid cultures and to the medium in the selection plates.

### Selection and genotyping of transformants

Approximately 100 transformants were grown to adult size on medium containing the hygromycin selection and 800 μM isoleucine, leucine and valine. One gemma was taken from each of 100 transformants and placed on medium supplemented with 800 μM branched chain amino acids, while a gemma from the same gemma cup was placed on medium without branched chain amino acids. Plants from which the gemma died or grew smaller on the medium lacking branched chain amino acids were then chosen as possible Mp*ALS* mutants and genotyped. Plants were genotyped using the Phire Plant Direct PCR kit (Thermo Fisher) using a 0.5 mm diameter piece of plant tissue and the manufacturer’s recommended 3-step cycling protocol. Primers used for genotyping can be found listed below.

### Sample preparation and RNA extraction

Total RNA was extracted from Tak-2 and target-site resistant mutants Mp*als*^*csP197L_7*^ and Mp*als*^*csP197L_10*^ across eight timepoints: 0h, 2h, 4h, 6h, 12h, 24h, 36h and 48h after transferring plants to herbicide-treated (5 nM) or untreated plates. The criteria for the selection of the herbicide dose was a dose causing a toxic effect without killing the plant, and was chosen based on the results from the dose response curve. There were two replicate samples per timepoint, and six plants per sample. Total RNA was extracted from 90 samples.

Gemmae were grown for 8 days on cellulose discs on untreated medium then moved onto chlorsulfuron-treated medium or untreated control plates. Plants were collected at the different timepoints after transfer onto chlorsulfuron and immediately frozen in liquid N2 and kept at −80°C. Samples were pulverised with a tissuelyser (Qiagen) and total RNA was extracted using RNeasy plant mini kits (Qiagen, Hilden, Germany). To remove DNA, RNA samples were DNase treated using the DNA-free kit by Ambion (Life Technologies) following manufacturer instructions. The quantity of RNA was measured on a Nanodrop ND-1000 spectrophotometer and RNA integrity was measured on an Agilent 2100 BioAnalyser (Agilent Technologies). The RNA integrity number (RIN) provided by the BioAnalyzer software ranges from 1 (a very degraded sample) to 10 (a mostly intact sample). Samples with RIN numbers ≥8 were suitable for cDNA library preparation following the Illumina RNA prep protocols (Welcome Trust Oxford Genomics Centre, Oxford). Illumina sequencing of 75 bp paired-end reads was conducted on the 90 cDNA libraries by an Illumina HiSeq4000. Samples were run over five lanes, and coverage was approximately 12 million reads per sample.

### Read pre-processing and mapping to reference transcriptome

Raw reads were returned as fastq.gz files. These were unzipped and their quality was viewed using FastQC v.0.11.7. Illumina adaptors and low-quality tails were quality trimmed using Trimmomatic-0.38 (62). Paired-end reads were interleaved, and ribosomal RNA was removed using Sortmerna-2.1 (63) and error corrected using BayesHammer (SPAdes-3.10.0) (64) with setting–only-error-correction. Clean reads were mapped to a reference transcriptome (JGI 3.1, obtained from https://marchantia.info/download/v31/) using Salmon (65).

### Differential expression analysis

The differences in expression of each gene per transcript between sample pairs were calculated using the DESeq 2 v.1.21.3 package (53) in the statistical software R. P-values were adjusted following the Benjamini-Hochberg procedure for controlling the false discovery rate (66). The criteria for differential gene expression was an adjusted p-value < 0.05 and a |log2(fold-change)| ≥1 (a fold-change of expression ≥ 2) between compared groups. Volcano plots were produced for each comparison. Expression differences were compared between untreated and treated Tak-2 2-48 hours after chlorsulfuron treatment.

### Gene Ontology term enrichment analysis

The VISEAGO R package (67) was used to identify GO terms significantly enriched in differentially expressed genes between treated and untreated plants, using the classic algorithm and Fisher’s exact test. The functional annotations obtained from Marpolbase (https://marchantia.info/download/MpTak_v6.1/) for the MpTakv6.1 transcriptome were used to create a custom GO database, because the transcriptome used for the RNA-Seq analysis (JGI 3.1) is poorly annotated with a large number of obsolete GO annotations. Takv6.1 annotations were associated with JGI 3.1 gene names using the gene correspondence table provided by Marpolbase. Multidimensional plots were generated showing clusters of GO terms into functional groups based on Wang’s semantic similarity distance and ward.D2 aggregation criterion.

### Real time quantitative PCR validation of candidate genes

The RNA-Seq experimental setup was repeated with fewer timepoints for real time quantitative PCR (RT-qPCR) validation. Plants were transferred to control or herbicide treated plates eight days after growth and collected 6, 12, 24, and 36 hours after treatment. Samples were collected and RNA was extracted and DNase treated as described above. 1 μg RNA was converted into cDNA using ProtoScript II RT (NEB), oligo(dT)s and Murine RNAse inhibitor (NEB) in a 20 μL reaction volume according to manufacturer instructions. Specific primers for the Mp*CYP* and Mp*GST* genes identified in the transcriptome analysis were designed using the Primer3Plus tool (https://primer3plus.com/cgi-bin/dev/primer3plus.cgi). Mp*APT* and Mp*ACT* were selected as reference genes (68). The list of primers used for RT-qPCR can be found below. Primer efficiencies were calculated for each primer pair by building a standard curve with a 1:5 dilution series of pooled cDNA (50, 10, 2, 0.4, 0.08 ng) and ranged between 1.8-2.2.

RT-qPCR experiments were performed with three biological replicates using a QuantStudio 7 cycler (Applied Biosystems). The 10 μl reaction mixture contained 5 μl 2X SYBR Green/ROX master mix (Applied Biosystems), 2 μl of 1:20 diluted cDNA, and 0.6 μl each of the forward and reverse primer, and 2.4 μl of RNAse free water. Each biological replicate was run in two technical replicates. The QuantStudio7 cycling method was as follows: 50°C for 2 min, 95°C for 10 min, then 40 cycles of 95°C for 15s and 60°C for 1 min. A melt curve analysis was conducted after PCR amplification to check primer specificity, with temperatures ranging from 60°C to 95°C increasing in 0.05°C s^−1^ increments. Relative expression was calculated using an adapted 2-(ΔΔCt) method that accounts for differences in primer efficiencies between the gene of interest and the reference gene(s), and incorporates multiple control genes (69). Technical replicate Ct values were averaged for each biological replicate. Each biological replicate Ct was then subtracted from the average Ct of the 3 untreated biological replicates to give ΔCt values. The calculated primer efficiency of each primer pair to the power of ΔCt gave the relative quantity (RQ). Expression of the gene of interest in each sample was calculated by dividing the RQ of that gene by the geometric mean of the RQ of the two reference genes in the same sample to give the normalised relative quantity (NRQ) (70). Significant differences between treated and untreated groups for each gene and timepoint were calculated by Student’s t-tests.

#### Vector construction for the generation of Mp*CYP* and Mp*GST* overexpression alleles

DNA was extracted from the Tak-2 wild-type line using a CTAB extraction method adapted from (59). Each of the up-regulated genes (Mp*GSTF7*, Mp*GSTF10*, Mp*GSTF15*, Mp*CYP813A5*, Mp*CYP822A1*) was PCR amplified from genomic Tak-2 DNA using CloneAmp HiFi Premix (Takara Bio). The primers used to amplify gene sequences are listed below. PCR products were run on 1% agarose gel and fragments of the correct size were extracted and purified using the GeneJET Gel Extraction Kit (ThermoFisher) following manufacturer instructions. The purified products were individually cloned into a directional entry vector pENTR D-TOPO (ThermoFisher) and transformed into *E. coli*. The plasmid was extracted and purified using a Minprep Kit (Qiagen), and then cloned by LR reaction into destination vector pMpGWB403 carrying the constitutive endogenous elongation factor alpha (Mp*EF1α)* promoter (71). The final vectors were transformed into *Agrobacterium tumefaciens* (GV3101) by electroporation. Agrobacterium-mediated transformation of *M. polymorpha* spores followed previously described methods (55).

### Real time quantitative PCR of Mp*GST* and Mp*CYP* overexpression transformants

RNA was extracted from 14-day old gemmalings from five independent lines transformed with an overexpression construct containing one of five chlorsulfuron-induced genes (Mp*GSTF7*, Mp*GSTF10*, Mp*GSTF15*, Mp*CYP813A5*, Mp*CYP822A1*). RNA was extracted using the RNeasy plant mini kit (Qiagen, Hilden, Germany). RNA samples were DNase treated using the DNA-free kit by Ambion (Life Technologies) according to manufacturer instructions. RNA was converted to cDNA as described in the transcriptome analysis. The primers used for RT-qPCR are listed below. RT-qPCR experiments were performed with three biological replicates using a QuantStudio7 cycler (Applied Biosystems). Each biological replicate sample was run in two technical replicates. The reaction mixture, cycling method and relative expression calculations were conducted as described in the transcriptome analysis.

### Herbicide sensitivity assay of Mp*GST* and Mp*CYP* overexpression lines

To characterise the sensitivity of Mp*GST-* and Mp*CYP-* over-expressing plants to chlorsulfuron, gemmae over-expressing one of the four candidate genes (Mp*GSTF7*, Mp*GSTF10*, Mp*CYP813A5* and Mp*CYP822A1*) and from an empty vector control were grown on medium supplemented with 30nM chlorsulfuron (non-lethal dose), 200 nM chlorsulfuron (lethal dose) or no herbicide. Significant differences in growth between each line and the empty vector control within each treatment were calculated by Student’s t-tests.

### Primers

The overhangs for CRISPR-Cas9 and Gateway cloning are underlined. Start and stop codons are in bold.

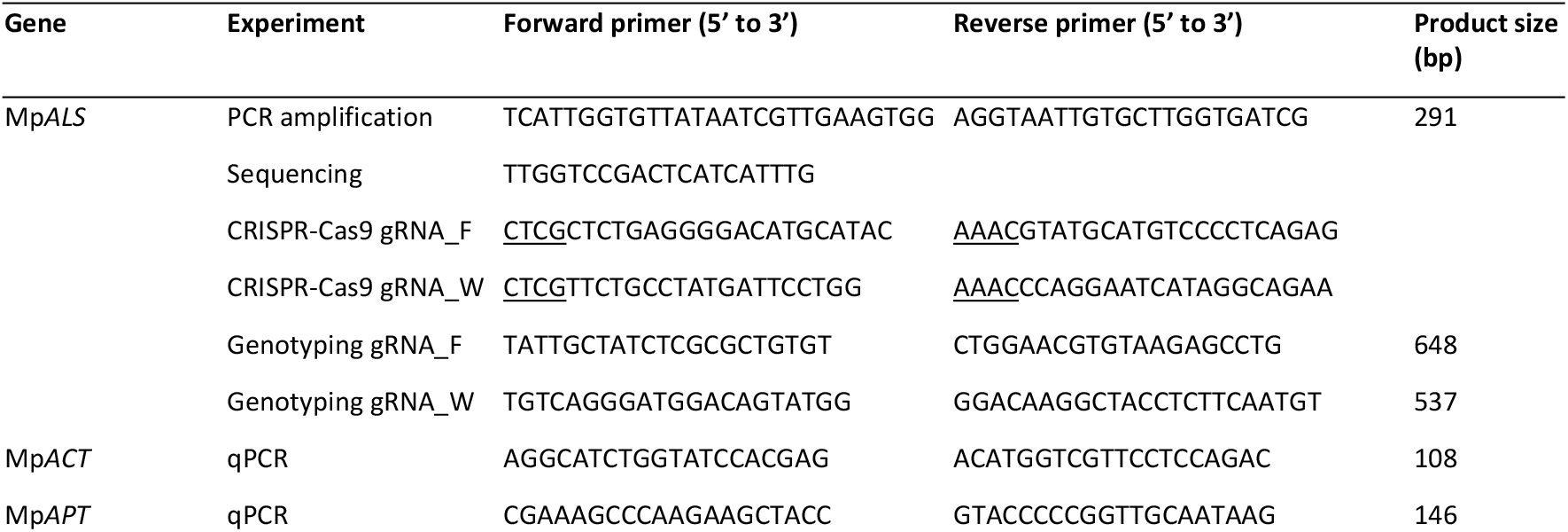
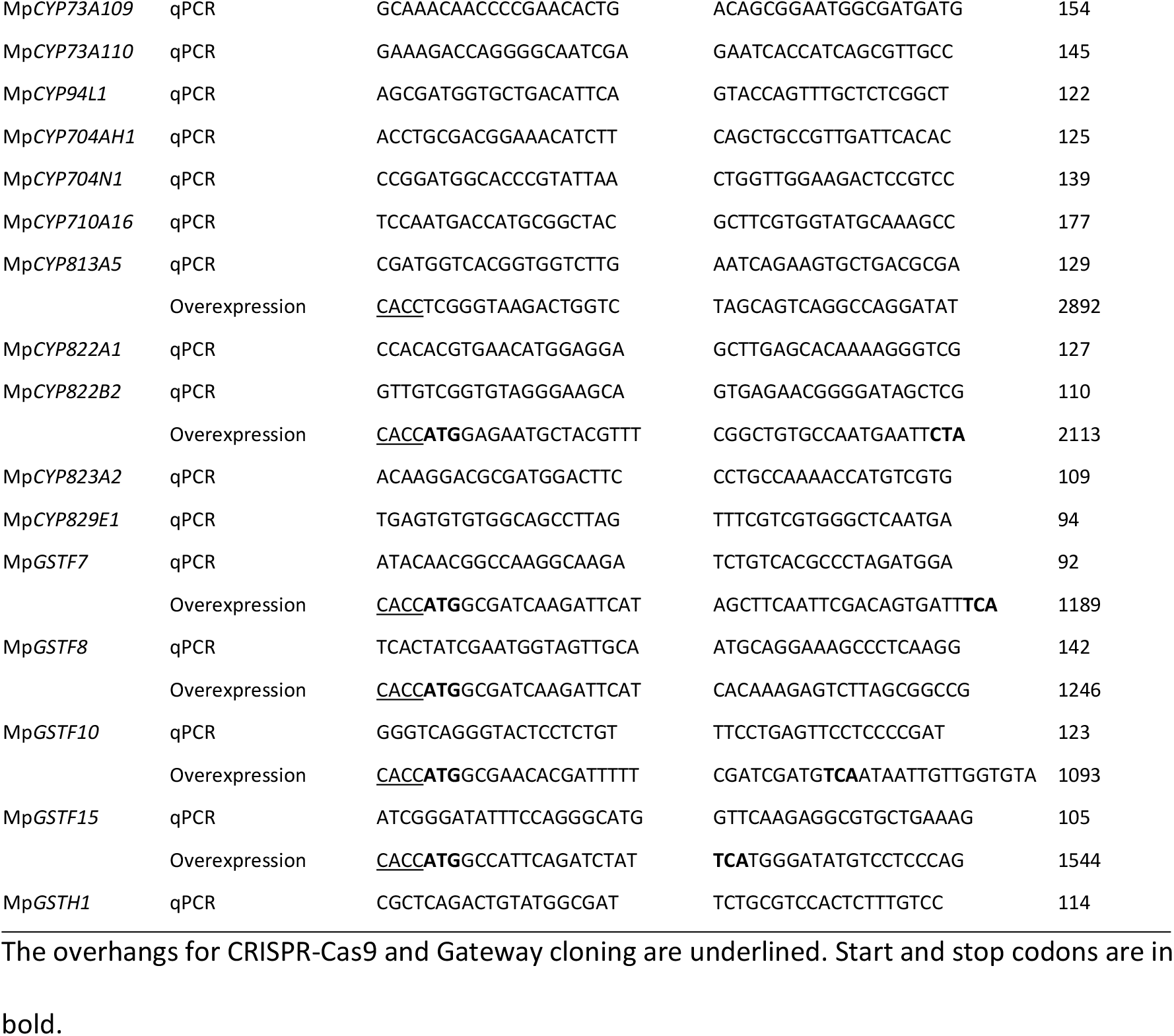

## Acknowledgments

We are grateful to Phil Poole, Laura Moody and Nick Kruger for suggestions. We are grateful to Simon Engledow for his help with processing the RNA-Seq samples. We thank Pippa Sinclair for laboratory assistance during the project. We thank Matt Watson for comments on the manuscript. We thank David Nelson from the University of Tennessee Health Science Center, who helped us with cytochrome P450 nomenclature.

## Supporting Information

**Table S1. List of metabolites detected in both RP and HILIC untargeted metabolomics analyses of chlorsulfuron-treated *M. polymorpha* (Fig 4C-E).**

Showing metabolite name, molecular weight (MW), retention time (RT), normalized peak area, Log_2_(fold change), *p*-value and adjusted *p*-value for each sample. C1-C6 are control samples. H1-H6 are herbicide-treated samples.

## References

1. Wang Y-M, Ong SS, Chai SC, Chen T. Role of CAR and PXR in xenobiotic sensing and metabolism. Expert Opin Drug Metab Toxicol. 2012 Jul;8(7):803.

2. Handschin C, Meyer UA. Induction of Drug Metabolism: The Role of Nuclear Receptors. Pharmacol Rev. 2003 Dec 1;55(4):649–73.

3. Morant M, Bak S, Møller BL, Werck-Reichhart D. Plant cytochromes P450: tools for pharmacology, plant protection and phytoremediation. Curr Opin Biotechnol. 2003 Apr 1;14(2):151–62.

4. Bai S, Liu W, Wang H, Zhao N, Jia S, Zou N, et al. Enhanced herbicide metabolism and metabolic resistance genes identified in tribenuron-methyl resistant Myosoton aquaticum L. J Agric Food Chem. 2018;66:9850–7.

5. Piasecki C, Yang Y, Benemann DP, Kremer FS, Galli V, Millwood RJ, et al. Transcriptomic analysis identifies new non-target site glyphosate-resistance genes in Conyza bonariensis. Plants. 2019 Jun 7;8(6):157.

6. Matzrafi M, Shaar-Moshe L, Rubin B, Peleg Z. Unraveling the Transcriptional Basis of Temperature-Dependent Pinoxaden Resistance in Brachypodium hybridum. Front Plant Sci. 2017 Jun 21;8:1064.

7. Iwakami S, Uchino A, Kataoka Y, Shibaike H, Watanabe H, Inamura T. Cytochrome P450 genes induced by bispyribac-sodium treatment in a multiple-herbicide-resistant biotype of Echinochloa phyllopogon. Pest Manag Sci. 2014 Apr 1;70(4):549–58.

8. Wang J, Chen J, Li X, Cui H. RNA-Seq transcriptome analysis to identify candidate genes involved in non-target site-based mesosulfuron-methyl resistance in Beckmannia syzigachne. Pestic Biochem Physiol. 2021 Jan 1;171:104738.

9. Leslie T, Baucom RS. De novo assembly and annotation of the transcriptome of the agricultural weed Ipomoea purpurea uncovers gene expression changes associated with herbicide resistance. G3 Genes, Genomes, Genet. 2014 Oct 1;4(10):2035–47.

10. Zhao N, Li W, Bai S, Guo W, Yuan G, Wang F, et al. Transcriptome profiling to identify genes involved in mesosulfuron-methyl resistance in Alopecurus aequalis. Front Plant Sci. 2017 Aug 9;8:1391.

11. Franco-Ortega S, Goldberg-Cavalleri A, Walker A, Brazier-Hicks M, Onkokesung N, Edwards R. Non-target site herbicide resistance is conferred by two distinct mechanisms in black-grass (Alopecurus myosuroides). Front Plant Sci. 2021 Mar 3;12:636652.

12. Liu W, Bai S, Zhao N, Jia S, Li W, Zhang L, et al. Non-target site-based resistance to tribenuronmethyl and essential involved genes in Myosoton aquaticum (L.). BMC Plant Biol. 2018 Oct 11;18(1):225.

13. Cabello-Hurtado F, Batard Y, Salaün JP, Durst F, Pinot F, Werck-Reichhart D. Cloning, expression in yeast, and functional characterization of CYP81B1, a plant cytochrome P450 that catalyzes in-chain hydroxylation of fatty acids. J Biol Chem. 1998 Mar 27;273(13):7260–7.

14. Han H, Yu Q, Beffa R, González S, Maiwald F, Wang J, et al. Cytochrome P450 CYP81A10v7 in Lolium rigidum confers metabolic resistance to herbicides across at least five modes of action. Plant J. 2021 Jan 27;105(1):79–92.

15. Gaines TA, Lorentz L, Figge A, Herrmann J, Maiwald F, Ott M-C, et al. RNA-Seq transcriptome analysis to identify genes involved in metabolism-based diclofop resistance in Lolium rigidum. Plant J. 2014 Jun;78(5):865–76.

16. Duhoux A, Carrère S, Gouzy J, Bonin L, Délye C. RNA-Seq analysis of rye-grass transcriptomic response to an herbicide inhibiting acetolactate-synthase identifies transcripts linked to non-target-site-based resistance. Plant Mol Biol. 2015 Mar 31;87(4–5):473–87.

17. Duhoux A, Carrère S, Duhoux A, Délye C. Transcriptional markers enable identification of ryegrass (Lolium sp.) plants with non-target-site-based resistance to herbicides inhibiting acetolactate-synthase. Plant Sci. 2017 Apr 1;257:22–36.

18. Pan L, Gao H, Xia W, Zhang T, Dong L. Establishing a herbicide-metabolizing enzyme library in Beckmannia syzigachne to identify genes associated with metabolic resistance. J Exp Bot. 2016 Mar 1;67(6):1745–57.

19. Salas-Perez RA, Saski CA, Noorai RE, Srivastava SK, Lawton-Rauh AL, Nichols RL, et al. RNA-Seq transcriptome analysis of Amaranthus palmeri with differential tolerance to glufosinate herbicide. PLoS One. 2018;13:1–33.

20. Thyssen GN, Naoumkina M, McCarty JC, Jenkins JN, Florane C, Li P, et al. The P450 gene CYP749A16 is required for tolerance to the sulfonylurea herbicide trifloxysulfuron sodium in cotton (Gossypium hirsutum L.). BMC Plant Biol. 2018 Dec 10;18(1):186.

21. Zhao N, Yan Y, Liu W, Wang J. Cytochrome P450 CYP709C56 metabolizing mesosulfuronmethyl confers herbicide resistance in Alopecurus aequalis. Cell Mol Life Sci. 2022 Apr 1;79(4):1–14.

22. Iwakami S, Endo M, Saika H, Okuno J, Nakamura N, Yokoyama M, et al. Cytochrome P450 CYP81A12 and CYP81A21 are associated with resistance to two acetolactate synthase inhibitors in Echinochloa phyllopogon. Plant Physiol. 2014 Jun 1;165(2):618–29.

23. Yanniccari M, Gigón R, Larsen A. Cytochrome P450 Herbicide Metabolism as the Main Mechanism of Cross-Resistance to ACCase- and ALS-Inhibitors in Lolium spp. Populations From Argentina: A Molecular Approach in Characterization and Detection. Front Plant Sci. 2020 Nov 16;11:1813.

24. Dimaano NG, Yamaguchi T, Fukunishi K, Tominaga T, Iwakami S. Functional characterization of cytochrome P450 CYP81A subfamily to disclose the pattern of cross-resistance in Echinochloa phyllopogon. Plant Mol Biol. 2020 Mar 1;102(4–5):403–16.

25. Iwakami S, Kamidate Y, Yamaguchi T, Ishizaka M, Endo M, Suda H, et al. CYP81A P450s are involved in concomitant cross-resistance to acetolactate synthase and acetyl-CoA carboxylase herbicides in Echinochloa phyllopogon. New Phytol. 2019 Mar 1;221(4):2112–22.

26. Ramel F, Sulmon C, Serra A-A, Gouesbet G, Couée I. Xenobiotic sensing and signalling in higher plants. J Exp Bot. 2012 Jun 28;63(11):3999–4014.

27. Alberto D, Serra AA, Sulmon C, Gouesbet G, Couée I. Herbicide-related signaling in plants reveals novel insights for herbicide use strategies, environmental risk assessment and global change assessment challenges. Vols. 569–570, Science of the Total Environment. Elsevier B.V.; 2016. p. 1618–28.

28. Ray TB. The mode of action of chlorsulfuron: A new herbicide for cereals. Pestic Biochem Physiol. 1982 Feb 1;17(1):10–7.

29. Ray TB. Site of Action of Chlorsulfuron. Plant Physiol. 1984 Jul;75(3):827–31.

30. Rhodes D, Hogan AL, Deal L, Jamieson GC, Haworth P. Amino Acid Metabolism of Lemna minor L. Plant Physiol. 1987 Jul 1;84(3):775–80.

31. Zabalza A, Orcaray L, Igal M, Schauer N, Fernie AR, Geigenberger P, et al. Unraveling the role of fermentation in the mode of action of acetolactate synthase inhibitors by metabolic profiling. J Plant Physiol. 2011 Sep 1;168(13):1568–75.

32. Scheel D, Casida JE. Sulfonylurea herbicides: Growth inhibition in soybean cell suspension cultures and in bacteria correlated with block in biosynthesis of valine, leucine, or isoleucine. Pestic Biochem Physiol. 1985 Jun 1;23(3):398–412.

33. Shaner DL, Singh BK. Phytotoxicity of Acetohydroxyacid Synthase Inhibitors Is Not Due to Accumulation of 2-Ketobutyrate and/or 2-Aminobutyrate. Plant Physiol. 1993 Dec 1;103(4):1221–6.

34. Huang T, Jander G. Abscisic acid-regulated protein degradation causes osmotic stress-induced accumulation of branched-chain amino acids in Arabidopsis thaliana. Planta. 2017 Oct 1;246(4):737–47.

35. Orcaray L, Igal M, Marino D, Zabalza A, Royuela M. The possible role of quinate in the mode of action of glyphosate and acetolactate synthase inhibitors. Pest Manag Sci. 2010 Mar 1;66(3):262–9.

36. Trenkamp S, Eckes P, Busch M, Fernie AR. Temporally resolved GC-MS-based metabolic profiling of herbicide treated plants treated reveals that changes in polar primary metabolites alone can distinguish herbicides of differing mode of action. Metabolomics. 2009 Dec 13;5(3):277–91.

37. Heap I. The International Survey of Herbicide Resistant Weeds [Internet]. 2021. Available from: www.weedscience.org

38. Tranel PJ, Wright TR. Resistance of weeds to ALS-inhibiting herbicides: what have we learned? Weed Sci. 2002;50:700–12.

39. Yu Q, Powles SB. Resistance to AHAS inhibitor herbicides: current understanding. Pest Manag Sci. 2014 Sep;70(9):1340–50.

40. McCourt JA, Pang SS, King-Scott J, Guddat LW, Duggleby RG. Herbicide-binding sites revealed in the structure of plant acetohydroxyacid synthase. Proc Natl Acad Sci. 2006;103(3):569–73.

41. Liu W, Yuan G, Du L, Guo W, Li L, Bi Y, et al. A novel Pro197Glu substitution in acetolactate synthase (ALS) confers broad-spectrum resistance across ALS inhibitors. Pestic Biochem Physiol. 2015 Jan 1;117:31–8.

42. Liu W, Bi Y, Li L, Yuan G, Du L, Wang J. Target-site basis for resistance to acetolactate synthase inhibitor in Water chickweed (Myosoton aquaticum L.). Pestic Biochem Physiol. 2013 Sep 1;107(1):50–4.

43. Rey-Caballero J, Menéndez J, Osuna MD, Salas M, Torra J. Target-site and non-target-site resistance mechanisms to ALS inhibiting herbicides in Papaver rhoeas. Pestic Biochem Physiol. 2017 May 1;138:57–65.

44. Yuan JS, Tranel PJ, Stewart CN. Non-target-site herbicide resistance: a family business. Trends Plant Sci. 2007 Jan 1;12(1):6–13.

45. Newby A, Altland JE, Gilliam CH, Wehtje G. Pre-emergence Liverwort Control in Nursery Containers. Horttechnology. 2007 Jan 1;17(4):496–500.

46. Sidhu MK, Lopez RG, Chaudhari S, Saha D. A Review of Common Liverwort Control Practices in Container Nurseries and Greenhouse Operations. Horttechnology. 2020 Aug 1;30(4):471–9.

47. Yang Q, Deng W, Li X, Yu Q, Bai L, Zheng M. Target-site and non-target-site based resistance to the herbicide tribenuron-methyl in flixweed (Descurainia sophia L.). BMC Genomics. 2016;17(1):17:551.

48. Rojano-Delgado AM, Portugal JM, Palma-Bautista C, Alcántara-de la Cruz R, Torra J, Alcántara E, et al. Target site as the main mechanism of resistance to imazamox in a Euphorbia heterophylla biotype. Sci Rep. 2019 Dec 1;9(1).

49. Casey A, Dolan L. Genes encoding cytochrome P450 monooxygenases and glutathione S-transferases associated with herbicide resistance evolved before the origin of land plants. bioRxiv. 2022 Aug 15;2022.08.12.503801.

50. Dimaano NG, Iwakami S. Cytochrome P450-mediated herbicide metabolism in plants: current understanding and prospects. Pest Manag Sci. 2021 Jan 31;77(1):22–32.

51. Siminszky B. Plant cytochrome P450-mediated herbicide metabolism. Vol. 5, Phytochemistry Reviews. 2006. p. 445–58.

52. Cummins I, Dixon DP, Freitag-Pohl S, Skipsey M, Edwards R. Multiple roles for plant glutathione transferases in xenobiotic detoxification. Drug Metab Rev. 2011;43(2):266–80.

53. Love MI, Huber W, Anders S. Moderated estimation of fold change and dispersion for RNA-seq data with DESeq2. Genome Biol. 2014;15(12):550.

54. Délye C, Duhoux A, Gardin JAC, Gouzy J, Carrère S. High conservation of the transcriptional response to acetolactate-synthase-inhibiting herbicides across plant species. Iannetta P, editor. Weed Res. 2018 Feb 1;58(1):2–7.

55. Ishizaki K, Chiyoda S, Yamato KT, Kohchi T. Agrobacterium-mediated transformation of the haploid liverwort Marchantia polymorpha L., an emerging model for plant biology. Plant Cell Physiol. 2008;49(7):1084–91.

56. Johnson CM, Stout PR, Broyer TC, Carlton AB. Comparative chlorine requirements of different plant species. Plant Soil. 1957;8:337–53.

57. Gamborg OL, Miller RA, Ojima K. Nutrient requirements of suspension cultures of soybean root cells. Exp Cell Res. 1968 Apr 1;50(1):151–8.

58. Chiyoda S, Ishizaki K, Kataoka H, Yamato KT, Kohchi T. Direct transformation of the liverwort Marchantia polymorpha L. by particle bombardment using immature thalli developing from spores. Plant Cell Rep. 2008 Sep 14;27(9):1467–73.

59. Porebski S, Bailey LG, Baum BR. Modification of a CTAB DNA extraction protocol for plants containing high polysaccharide and polyphenol components. Vol. 15, Plant Molecular Biology Reporter. International Society for Plant Molecular Biology; 1997. p. 8–15.

60. Sugano SS, Shirakawa M, Takagi J, Matsuda Y, Shimada T, Hara-Nishimura I, et al. CRISPR/Cas9-Mediated Targeted Mutagenesis in the Liverwort Marchantia polymorpha L. Plant Cell Physiol. 2014 Mar 1;55(3):475–81.

61. Lei Y, Lu L, Liu HY, Li S, Xing F, Chen LL. CRISPR-P: A web tool for synthetic single-guide RNA design of CRISPR-system in plants. Vol. 7, Molecular Plant. Cell Press; 2014. p. 1494–6.

62. Bolger AM, Lohse M, Usadel B. Trimmomatic: a flexible trimmer for Illumina sequence data. Bioinformatics. 2014 Aug 1;30(15):2114–20.

63. Kopylova E, Noé L, Touzet H. SortMeRNA: fast and accurate filtering of ribosomal RNAs in metatranscriptomic data. Bioinformatics. 2012 Dec 1;28(24):3211–7.

64. Nikolenko SI, Korobeynikov AI, Alekseyev MA. BayesHammer: Bayesian clustering for error correction in single-cell sequencing. BMC Genomics. 2013 Jan 21;14(Suppl 1):S7.

65. Patro R, Duggal G, Love MI, Irizarry RA, Kingsford C. Salmon provides fast and bias-aware quantification of transcript expression. Nat Methods. 2017 Apr 6;14(4):417–9.

66. Benjamini Y, Hochberg Y. Controlling the false discovery rate: a practical and powerful approach to multiple testing. Vol. 57, Journal of the Royal Statistical Society. 1995.

67. Brionne A, Juanchich A, Hennequet-Antier C. ViSEAGO: A Bioconductor package for clustering biological functions using Gene Ontology and semantic similarity. BioData Min. 2019 Aug 6;12(1).

68. Saint-Marcoux D, Proust H, Dolan L, Langdale JA. Identification of Reference Genes for Real-Time Quantitative PCR Experiments in the Liverwort Marchantia polymorpha. Margis R, editor. PLoS One. 2015 Mar 23;10(3):e0118678.

69. Hellemans J, Mortier G, De Paepe A, Speleman F, Vandesompele J. qBase relative quantification framework and software for management and automated analysis of real-time quantitative PCR data. Genome Biol. 2008 Feb 9;8(2):R19.

70. Vandesompele J, De Preter K, Pattyn F, Poppe B, Van Roy N, De Paepe A, et al. Accurate normalization of real-time quantitative RT-PCR data by geometric averaging of multiple internal control genes. Genome Biol. 2002;3(7):research0034.1.

71. Ishizaki K, Nishihama R, Ueda M, Inoue K, Ishida S, Nishimura Y, et al. Development of Gateway Binary Vector Series with Four Different Selection Markers for the Liverwort Marchantia polymorpha. Ezura H, editor. PLoS One. 2015 Sep 25;10(9):e0138876.

